# A dominant-negative SOX18 mutant disrupts multiple regulatory layers essential to transcription factor activity

**DOI:** 10.1101/2020.11.11.378968

**Authors:** Alex McCann, Jieqiong Lou, Mehdi Moustaqil, Ailisa Blum, Frank Fontaine, Hui Liu, Winnie Luu, Peter Koopman, Emma Sierecki, Yann Gambin, Frédéric A. Meunier, Zhe Liu, Elizabeth Hinde, Mathias Francois

## Abstract

Few genetically dominant mutations involved in human disease have been fully explained at the molecular level. In cases where the mutant gene encodes a transcription factor, the dominant-negative mode of action of the mutant protein is particularly poorly understood. Here, we studied the genome-wide mechanism underlying a dominant-negative form of the SOX18 transcription factor (SOX18^RaOp^) responsible for both the classical mouse mutant Ragged opossum and the human genetic disorder Hypotrichosis-Lymphedema-Telangiectasia-Renal Syndrome. Combing three single-molecule imaging assays in living cells, we found that SOX18^RaOp^ disrupts the system through an accumulation of molecular interferences which impair several functional properties of the wild-type SOX18 protein, including its chromatin-binding dynamics. The dominant-negative effect is further amplified by recruiting the interactome of its wild-type counterpart, which perturbs regulatory nodes such as SOX7 and MEF2C. Our findings explain in unprecedented detail the multi-layered process that underpins the molecular etiology of dominant-negative transcription factor function.

## Introduction

Embryonic development is dependent upon the activity of transcription factor (TF) complexes, which assemble on the chromatin in a finely orchestrated temporal and spatial sequence to coordinate the expression of specific gene programs (*1, 2*). A current challenge in the study of human genetic disease is to understand how perturbed TF function leads to the phenotypic spectrum at the molecular level. For recessive disorders, loss or impairment of TF function provides a ready explanation. However, dominant disorders have been more difficult to pin down, but are ascribed to dominant-negative TF activity, whereby a mutant protein interferes with the functionality of its wild-type counterpart. Concepts such as neomorphism (where the mutated gene product takes on novel functions) and antimorphism (where the mutated gene product antagonizes the wild-type gene product) are commonly evoked (*3*). Despite this, it is currently not clear whether or how these concepts manifest at the molecular level; nor is it clear whether these are the only possible modes of action of dominant inheritance. Several classes of dominant-negative TF mutations have been described, including those causing truncation, deletion or alteration of either the DNA-binding domain or another functional domain (*4*). However, the key to understanding how these mutant TFs act in a dominant-negative fashion lies in discovering not only how the mutant protein lacks the function of the normal protein, but also how is actively interferes with the function of the normal protein (*5*).

TFs function primarily as heterodimers or homodimers. Where a dominant-negative TF is expressed, it has been proposed that the overall protein functionality of TFs that form homodimers will be only 25% of the wild-type, since of the four potential homodimer configurations (wt/wt, wt/mut, mut/wt, mut/mut) only one (wt/wt) is functional, with the mutant form effectively poisoning the other complexes (*5*). Moreover, over-expression of a dominant-negative allele would amplify the observed negative effects, bringing functionality to below 25% (*5*). These scenarios potentially explain why dominant-negative mutations are usually more severe than recessive, loss-of-function mutations that would be predicted to reduce TF activity to ∼50% of the normal level.

Dominant-negative mutations causing genetic disorders have been observed to occur in genes encoding a number of TF in the SOX family (*6–10*). SOX TFs are key molecular switches of cell fate in numerous tissue types during embryogenesis (*11*). These include SOX8, 9 and 10 in which dominant-negative mutations trigger sex determination disorders (*8*), skeletal defects (*12*), and neural crest dysfunction (*10*) respectively.

Here, we have studied a dominant-negative mutation affecting the TF SOX18. Over the last 70 years, the study of SOX18 and its mutants in mice, humans and zebrafish have yielded profound insights into the regulation of vascular, lymphatic and hair follicle development (*7, 13–18*), and into the concepts of allelic series, genetic redundancy and the mode of action of genetic modifiers (*19, 20*). Despite an expression pattern suggesting an important role in vascular and hair follicle development (*15*), early attempts to understand the developmental role of SOX18 through the generation of *Sox18*-null mice were confounded by the surprisingly mild phenotype of normal vascular development with only mild hair follicle anomalies (*21*). SOX18 is co-expressed with closely related ‘SOXF’ subfamily members SOX7 and SOX17 in endothelial cells (*19, 20, 22*), suggesting that the lack of phenotype of *Sox18*-null mice is due to functional redundancy between the three TFs.

This phenotype contrasts with that of a classical mouse mutant, Ragged, found to result from nonsense mutations in Sox18 (*15*). Discovered in the 1950s (*13, 14*), four Ragged alleles were described, the most severe of which is known as Ragged opossum (SOX18^RaOp^) (*23, 24*). Inheritance of one Sox18^RaOp^ allele is sufficient to yield a thin, ragged coat and vascular leakage. The presence of two mutant alleles together causes death *in utero* due to lethal vascular dysfunction and/or lymphedema (*25*). The dramatically different phenotypes of *Sox18*-null and SOX18^RaOp^ mice suggested a dominant-negative mechanism of the latter, whereby SOX18^RaOp^ interferes with the function of SOX18 and also SOX7 and SOX17. While there has been some progress in elucidating the genetic pathways downstream of SOX18 (*16, 26–28*), it remains unclear how the mutant SOX18^RaOp^ protein interferes with wild-type SOX18 function to perturb downstream gene expression.

Analogous to SOX18^RaOp^ mice, dominant mutations in SOX18 in humans cause the rare congenital disorder Hypotrichosis-Lymphedema-Telangiectasia-Renal Syndrome (HLTRS) (*6, 7*). Patients diagnosed with HLTRS exhibit prominent hair follicle and vascular defects; hair follicles are sparse or absent, lymphatic vessels leak causing swollen limbs, and various cardiovascular defects are present, including those that can cause renal failure. An unexplained etiological component of HLTRS is that SOX18 mutations, akin to mouse, occur in allelic series which underpin the severity of the syndrome with defects ranging from mild to lethal (*29*).

An understanding of how dominant-negative SOX18 proteins give rise to a range of phenotypic outcomes in SOX18^RaOp^ mice and HLTRS children rests on detailed knowledge of the mode of action of the wild-type SOX18 TF. Here, we apply a suite of molecular imaging assays to visualise SOX18 nuclear dynamics and analyse its search pattern on the chromatin. We use this pipeline to quantify dominant-negative effects of mutant SOX18 proteins beyond simple genetic configuration and level of allele expression. We show that altered biophysical parameters such as nuclear concentration, diffusion, oligomeric state, chromatin-binding dynamics, chromatin-binding affinity, and protein stability unbalance the regulatory network in favour of the mutant protein. This study defines novel mechanisms of interference that underlie dominant-negative TF action, providing new insights into the mechanisms that underpin dominant human genetic disorders.

## Results

In order to analyze the search pattern mechanism of SOX18 and its dominant-negative mutant counterpart, we took advantage of self-labelling Halo-tag technology (*30, 31*) and used it in combination with three single-molecule resolution imaging techniques – single molecule tracking (SMT) (*1, 2, 32–36*) to measure the overall chromatin-binding dynamics, number and brightness (N&B) (*37–39*) analysis to obtain the oligomeric distribution and cross-RICS (cRICS) (*40*) analysis using two spectrally distinct Halo-tag dyes (JF549 and JF646) to validate homodimer formation and to obtain the chromatin-bound fraction of homodimers. An overview describing the steps involved in, and biological information obtained from using each of these imaging techniques, is described in Fig. S1, which is a supporting figure to Fig. 1 (Fig. S1 to Fig. 1).

**Fig. 1.**
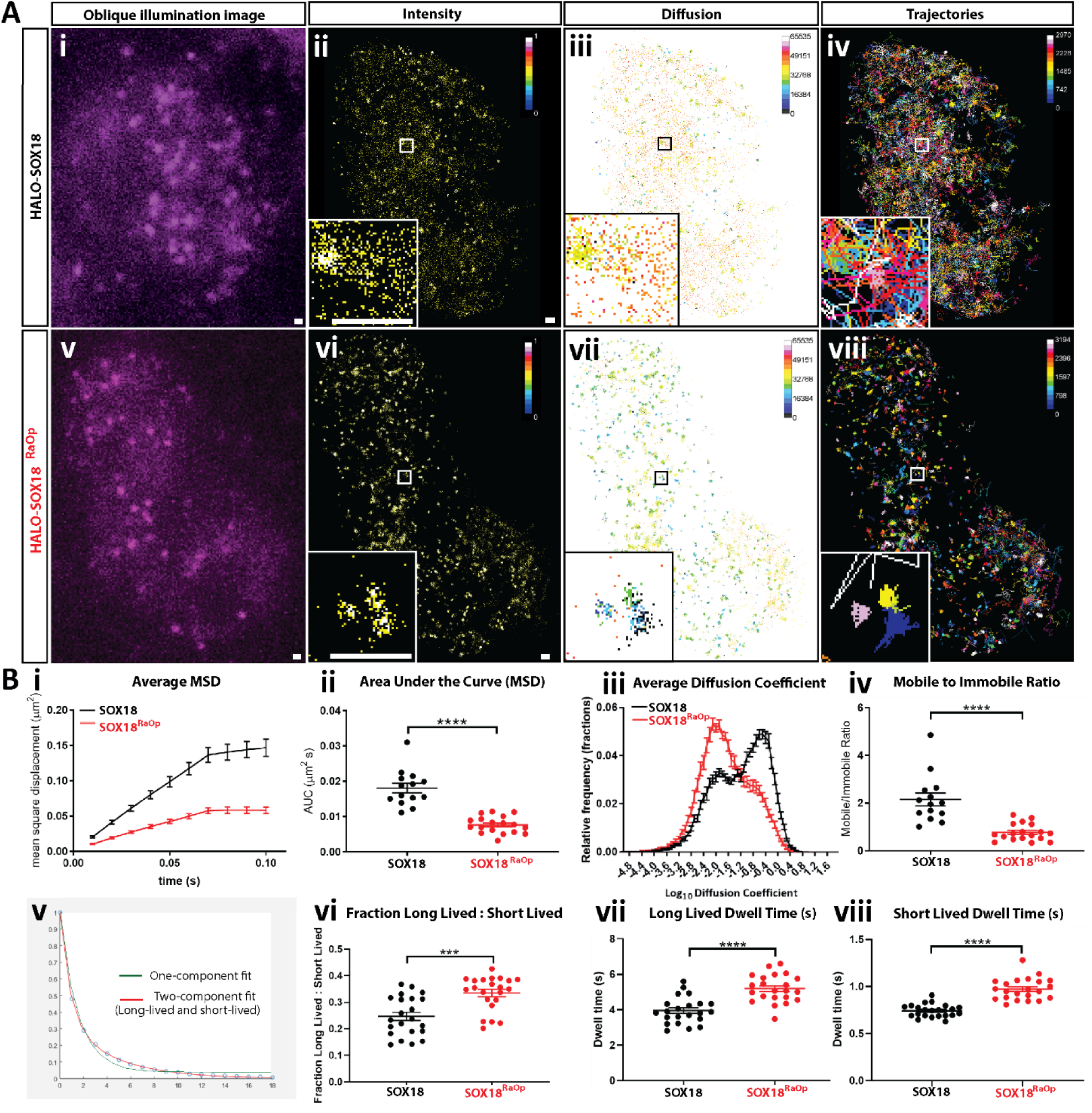
The SOX18^RaOp^ dominant-negative mutant protein displays impaired chromatin-binding dynamics. (**A**) (**i and v**) snapshot from oblique illumination live imaging. Heat maps: **(ii and vi)** fluorescence intensity (color code: white = highest intensity, black = lowest intensity), **(iii and vii)** diffusion coefficient (color code: warmer colors = higher mobilities, cooler colors = lower mobilities) and (**iv and viii)** trajectory maps (color code: based on trajectory frame). Scale bar, 0.5 µm. Example image number of trajectories: HALO-SOX18 = 2970, HALO-SOX18^RaOp^ = 3194. Average number of trajectories: HALO-SOX18 = 2453, HALO-SOX18^RaOp^ = 2215. (**B**) Quantification of the dynamics of HALO-SOX18 (black) and HALO-SOX18^RaOp^ (red). **Top row:** (**i**) the average mean square displacement (MSD; μm^2^s), (**ii**) the area under the curve of the average MSD for each cell (μm^2^s), (**iii**) the diffusion coefficient histogram for all cells (μm^2^s-1) and (**iv**) the mobile to immobile ratio for each cell. Threshold to classify mobile and immobile molecules is Log_10_D = -1.5. Values for the mean ± s.e.m. are shown. n = 14 for HALO-SOX18 and n = 18 for HALO-SOX18^RaOp^ (N = 3). t-test (two-tailed, unpaired). **** P<0.0001. **Bottom row:** (**v**) a two-component-fit example for HALO-SOX18; (**vi**) fraction of long-lived to short-lived immobile events and dwell times of (**vii**) long-lived and (**viii**) short-lived immobile events (s). Values for the mean ± s.e.m. are shown. n = 22 for HALO-SOX18 and n = 22 for HALO-SOX18^RaOp^ (N = 3). Mann Whitney U-test for slow tracking (two-tailed, unpaired). *** P<0.001, **** P<0.0001.

Of note, SOX18 is a key regulator of blood vessel development, and as a result is highly expressed in blood vascular endothelial cells. In support of this, CAGE data from the FANTOM5 consortium shows that SOX18 is the most enriched TF in Human Umbilical Vascular Endothelial Cells (HUVECs) (*41*). Here, we intend to assess the mode of action of a dominant-negative SOX18 mutant (SOX18^RaOp^) in the absence of wild-type SOX18. For this reason, we have performed SMT, N&B and cRICS experiments in HeLa cells which have no detectable endogenous expression of SOX18 (Fig. S2A to Fig. 1).

### The SOX18^RaOp^ dominant-negative mutant form is a potent transcriptional repressor

Here, we set out to investigate the molecular mode of action of disease-causing mutations that give rise to a dominant-negative SOX18 protein. Such a truncated protein is still able to bind to DNA but fails to activate gene transcription (*42*). The Ragged mouse model exhibits natural mutations in the SOX18 gene – an allelic series of mutations associated with a broad range of phenotypic outcomes (from mild to severe vascular, renal and hair follicles defects). The most severe Ragged mutant is known as *opossum* (SOX18^RaOp^), characterized by a point deletion within the C-terminal transactivation domain causing a frameshift that scrambles the rest of the transactivation domain before resulting in a premature stop codon (Fig. S2B to Fig. 1). Due to the nature of its mutation, SOX18^RaOp^ is truncated and forms a shorter version of SOX18. To ensure that this size difference does not significantly affect the protein level by altering the degradation rate, we assessed the concentration of HALO-SOX18 and HALO-SOX18^RaOp^ at the whole cell and nuclear level (Fig. S2C to Fig. 1). By comparing whole cell lysates, we found that HALO-SOX18 and HALO-SOX18^RaOp^ are expressed at comparable levels, and by comparing nuclear lysates we found that more HALO-SOX18^RaOp^ accumulates in the nucleus. This shows that in addition to allele overexpression reducing protein functionality to below 25 % as described by Veitia (*5*), some dominant-negative mutations increase the intranuclear concentration of a TF, therefore amplifying its deleterious potential.

To explore the molecular mode of action of SOX18^RaOp^ during gene transactivation, we performed a luciferase assay using a synthetic VCAM-1 promoter fragment as a readout for SOX18 transcriptional activity (*43*) (Fig. S2D to Fig. 1). HALO-SOX18 efficiently activated VCAM-1 promoter activity, whereas SOX18^RaOp^ failed to do so. Further, SOX18^RaOp^ prevented HALO-SOX18 from transactivating the VCAM-1 promoter fragment. These results validate that the addition of the HALO-tag does not compromise SOX18 transcriptional activity nor does it prevent the dominant-negative mode of action of SOX18^RaOp^. The addition of an excess of HALO-SOX18 construct in a dose-dependent manner to outcompete the repressive effect of SOX18^RaOp^ failed to rescue the lack of transcriptional activity caused by the mutant protein. Strikingly, even at a ratio of 30:1 HALO-SOX18 to SOX18^RaOp^, the transcriptional activity was not restored, suggesting that gene dose response of the wild-type allele is not sufficient to compensate the dominant-negative mechanism of the mutant protein. This is in accordance with a dominant-negative phenotype, and what has been previously mathematically modelled by Veitia (*5*).

Previous studies have used single molecule tracking (SMT) to uncover functional aspects of the search patterns of TFs, and by doing so have shown that changes in their chromatin-binding behavior reflects changes in their gene target selection and activity (*1, 2, 33–36*). Here, we hypothesized that altered chromatin-binding dynamics may form an important part of the dominant-negative mode of action of SOX18^RaOp^. To explore this we generated a HALO-SOX18^RaOp^ construct using HALO-SOX18 as a backbone, transiently transfected HeLa cells with either HALO-SOX18 or HALO-SOX18^RaOp^, and performed fast (20ms acquisition, 6000 frames) and slow (500ms acquisition, 500 frames) SMT (Fig. 1A and 1B, Videos S1 and S2). These two different acquisition speeds provide us with information on the trajectories of the unbound diffusing and immobile chromatin-bound states (fast acquisition), and the different types of dwell times on the chromatin (slow acquisition).

Focusing on the chromatin-binding dynamics in the wild-type scenario, we found that an average of 32 % of SOX18 molecules are immobile, with the rest being mobile and diffusing (Fig. 1B). Of this immobile fraction, by comparing one and two-component fit models we identify that there are at least two types of immobile populations with different dwell times, which is in accordance with what has been reported previously for SOX2 (*2*) and other TFs (*2, 33–35, 44*). Here, we found that SOX18 had average long-lived dwell times of 3.94 s (Fig. 1B) and average short-lived dwell times of 0.74 s, in accordance with previously reported long-lived dwell times that typically last a few seconds (∼5-14.6 s) and short-lived dwell times that are typically 1 second or less (∼0.03–1.85 s) (*2, 33, 34, 44*). Additionally, we found that long-lived binding events accounted for one quarter of SOX18 immobile events. Previous SMT studies have demonstrated that these short-lived and long-lived dwell times are due to interactions with non-specific random and specific target chromatin sites respectively, notably via the use of DNA-binding and homodimerization mutants (*2, 34*).

When comparing the chromatin-binding dynamics of SOX18 to SOX18^RaOp^, we observed a significant difference in the search pattern of HALO-SOX18 and HALO-SOX18^RaOp^ already at the level of the raw data used for fast tracking analysis (Fig. 1A). An example of this highlighted in Video S1. HALO-SOX18^RaOp^ appears to be a lot less mobile than its wild-type counterpart, with what appears to be more immobile chromatin binding events, which remain immobile for longer (Video S1, Fig. 1A). The intensity and diffusion coefficient heatmaps show two main types of TF behaviors – immobile chromatin-binding events represented by distinct higher intensity regions associated with lower diffusion coefficients, and scattered between these, mobile diffusion events represented by lower intensity regions associated with higher diffusion coefficients (Fig. 1A). Based on this readout, SOX18^RaOp^ shows an overall lower mobility than SOX18, with more immobile chromatin-binding events and less diffusional events. In support of this, the trajectory maps show that SOX18^RaOp^ appears to have more trajectories contained within small areas suggesting greater immobility. By contrast, SOX18 appears to have more trajectories that explore less restricted areas, suggesting a higher diffusive behavior.

By quantifying the trajectories of HALO-SOX18 and HALO-SOX18^RaOp^ obtained via fast tracking SMT (Fig. 1B), we observed that overall SOX18^RaOp^ has a significantly lower mobility. This is based on its lower average mean square displacement (MSD), and its significantly higher immobile fraction represented by a higher peak in the diffusion coefficient histogram. The average MSDs for all trajectory types (mobile and immobile) shown in Fig. 1B were separated into average MSDs for immobile (Fig. S3A to Fig. 1) and mobile trajectories (Fig. S3B to Fig. 1). Comparing the average MSDs for immobile and mobile trajectories for HALO-SOX18 and HALO-SOX18^RaOP^ shows that while the mobility of HALO-SOX18^RaOp^ decreases in both immobile and mobile fractions, the mobile diffusing fraction is affected the most. Diffusion coefficient histograms for all cells are shown in Fig. S3C and S3D to Fig. 1. Despite heterogeneity in the diffusion coefficient histograms across cells, a clear trend can be observed where HALO-SOX18^RaOp^ shifts towards the immobile fraction.

By comparing the fraction of long-lived to short-lived immobile events obtained by slow tracking, and how long they occurred for, we found that SOX18^RaOp^ had a higher fraction of long-lived immobile events, and both long-lived and short-lived immobile events were longer for SOX18^RaOp^ than SOX18 (Fig. 1B). To assess whether a change in chromatin-binding stability may play a role in the difference in behavior observed for SOX18^RaOp^, we deleted the first alpha helix (AH1) of the HMG DNA-binding domain (HALO-SOX18^AH1^; Fig. S4 to Fig. 1). This mutant is still capable of binding to DNA via alpha helix 2 (AH2) but with lower affinity. As anticipated, decreasing the DNA-binding stability of SOX18 produced the opposite behavior observed for HALO-SOX18^RaOp,^ with SOX18^AH1^ displaying an increased MSD, lower chromatin-bound and long-lived binding fractions, and shorter dwell times. Of note, when SMT analysis was performed, only cells expressing a sufficient amount of HALO-SOX18^AH1^ in the nucleus were included, with cells that had less than 1000 trajectories excluded. Taken together these results indicate that one of the hallmarks of the non-functional SOX18^RaOp^ TF is a significant increase in chromatin-binding stability, which may also explain the enhanced nuclear concentration observed for SOX18^RaOp^ in Fig. S2A to Fig. 1.

### SOX18^RaOp^ mutant protein derails the chromatin-binding dynamics of SOX18

The dominant-negative form of SOX18 is embryonic lethal when homozygous, leaving only heterozygous individuals to survive this condition (*7, 45*). This implies that in the case of bi-allelic expression in a heterozygous scenario both SOX18 and SOX18^RaOp^ co-exist in the same cells at the same time. In order to assess the direct interference of SOX18^RaOp^ on SOX18 activity we next set out to measure the chromatin-binding dynamics of the wild-type protein in presence of the mutant protein.

To achieve this, SMT analysis was performed using transiently co-transfected HeLa cells with HALO-SOX18 in a 3:1 or 1:1 ratio with either untagged SOX18^RaOp^, or untagged SOX18 as a protein expression control (Fig. 2A and 2B). Example SMT videos comparing each of these conditions can be found in Videos S3 and S4. By quantifying and comparing the trajectories for HALO-SOX18:SOX18 (3:1) and HALO-SOX18:SOX18 (1:1), we found that there were no significant differences in the chromatin-binding dynamics of HALO-SOX18 despite increasing the amount of untagged protein. By using HALO-SOX18:SOX18 (3:1) and (1:1) as a point of comparison to HALO-SOX18:SOX18^RaOp^ (3:1) and (1:1) respectively, we found a significant difference in the behavior of HALO-SOX18 upon the addition of untagged-SOX18^RaOp^. By looking at the intensity, diffusion coefficient and trajectory maps, we found that HALO-SOX18 upon the addition of untagged- SOX18^RaOp^ behaves in a similar fashion to what was previously observed for HALO-SOX18^RaOp^, with HALO-SOX18 trajectories now having more distinct high intensity foci associated with lower diffusion coefficients and exploring less area (Fig. 2A). Examples of new behaviors are shown within the insets. These differences in behavior indicates that SOX18^RaOp^ greatly alters the search pattern of SOX18.

**Fig. 2.**
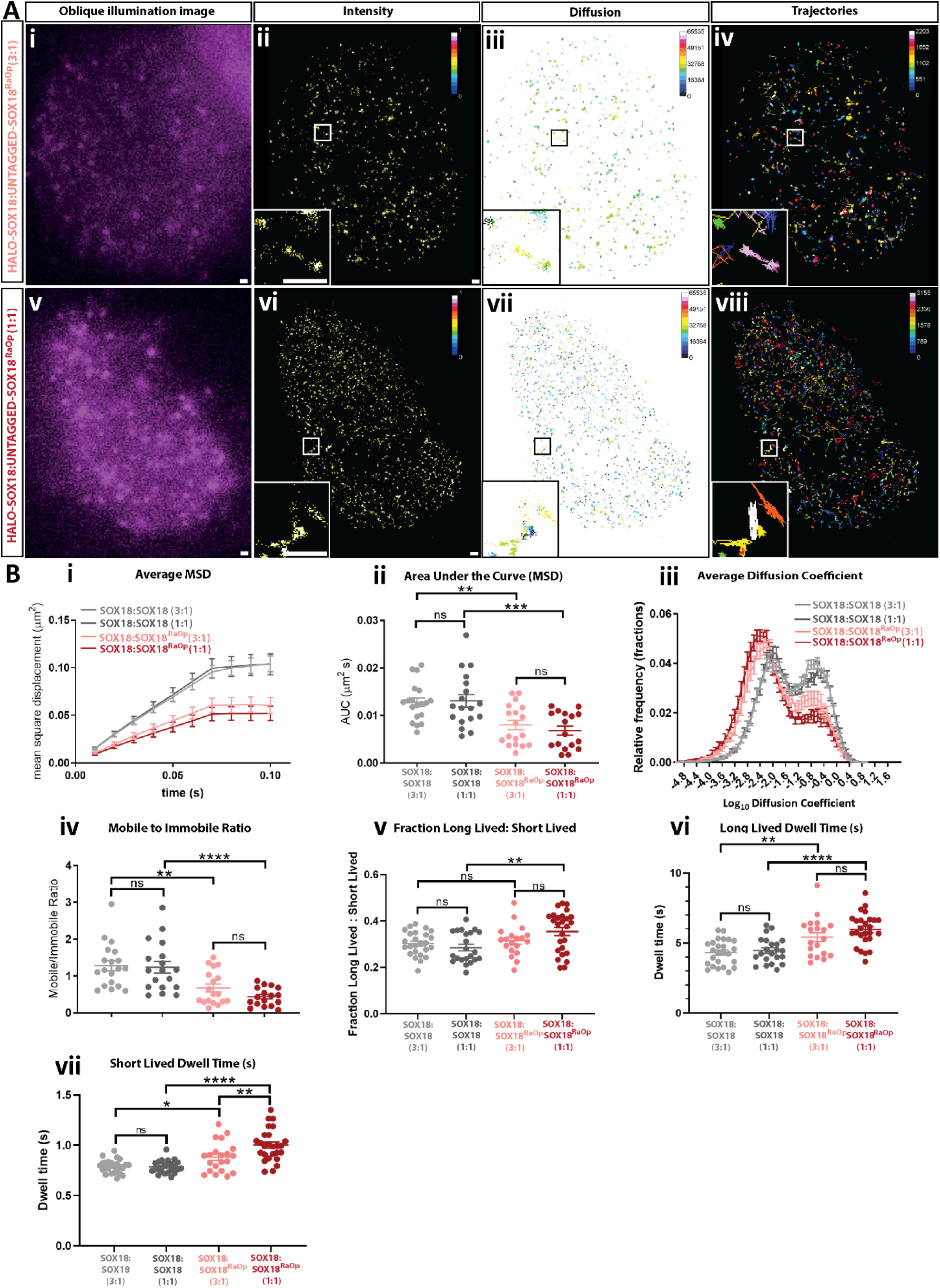
The SOX18^RaOp^ dominant-negative mutant protein directly interferes with the chromatin-binding dynamics of SOX18. (**A**) (**i and v**) snapshot from oblique illumination live imaging. Heat maps: **(ii and vi)** fluorescence intensity (color code: white = highest intensity, black = lowest intensity), **(iii and vii)** diffusion coefficient (color code: warmer colors = higher mobilities, cooler colors = lower mobilities) and (**iv and viii)** trajectory maps (color code: based on trajectory frame). Scale bar, 0.5 µm. Example image number of trajectories: HALO-SOX18-untagged-SOX18^RaOp^ (3:1) = 2203, HALO-SOX18:untagged-SOX18^RaOp^ (1:1) = 3155. Average number of trajectories: HALO-SOX18:untagged-SOX18 (3:1) = 2596, HALO-SOX18:untagged-SOX18 (1:1) = 1887, HALO-SOX18-untagged-SOX18^RaOp^ (3:1) = 1530, HALO-SOX18:untagged-SOX18^RaOp^ (1:1) = 1832. (**B**) Quantification of the dynamics of HALO-SOX18 with untagged SOX18 in a 3:1 ratio (light grey) or in a 1:1 ratio (dark grey) or with untagged SOX18^RaOp^ in a 3:1 ratio (light red) or 1:1 ratio (dark red). (**i**) average mean square displacement (MSD; μm^2^s), (**ii**) area under the curve (μm^2^s), (**iii**) average diffusion coefficient (μm^2^/s) and (**iv**) mobile to immobile ratio. Threshold to classify mobile and immobile molecules is Log_10_D = -1.5. Values for the mean ± s.e.m. are shown. n = 19 for HALO-SOX18:SOX18 (3:1), n = 18 for HALO-SOX18:SOX18 (1:1), n = 18 for HALO-SOX18:SOX18^RaOp^ (3:1) and n = 17 for HALO-SOX18:SOX18^RaOp^ (1:1) (N = 3). t-test (two-tailed, unpaired). ** P<0.01, *** P<0.001, **** P<0.0001, ns = non-significant (P>0.05). (**v**) fraction of long-lived to short-lived immobile events, and dwell times of **(vi)** long-lived and (**vii**) short-lived immobile events. Values for the mean ± s.e.m. are shown. n = 25 for HALO-SOX18:SOX18 (3:1), n = 22 for HALO-SOX18:SOX18 (1:1), n = 19 for HALO-SOX18:SOX18^RaOp^ (3:1) and n = 26 for HALO-SOX18:SOX18^RaOp^ (1:1) (N = 3). Mann Whitney U-test (two-tailed, unpaired). * P<0.05, ** P<0.01, **** P<0.0001, ns = non-significant (P>0.05).

Quantitation of HALO-SOX18 trajectories in the presence of SOX18^RaOp^ revealed that the addition of the dominant-negative protein decreased the overall mobility of SOX18 (Fig. 2B). Further, it also increased the immobilized fraction of the total SOX18 population, increased the fraction of immobile events that occurred for a long period of time, and extended the length of time of both long-lived and short-lived immobile events. Based on these observations, SOX18^RaOp^ appears to poison the wild-type TF with a SOX18^RaOp^-like molecular behavior.

### SOX18^RaOp^ recruits SOXF factors to form nonfunctional complexes

The SOX18 TF has been reported to act via an array of multiple protein-protein interactions (PPIs) (*42*), thus as a broad spectrum mechanism for interference, we assessed whether SOX18^RaOp^ may have the potential to affect SOX18 PPIs. At first, we took advantage of the previous identification of the SOX18 interactome (*42*) (Fig. 3A). We assessed whether different naturally occurring recessive mutations and dominant-negative mutants mimicking those reported in Human (*6, 7*) were able to retain their interaction with SOX18 protein partners by performing a protein-protein interaction assay using AlphaScreen technology. Here we show two main types of mutations – recessive mutations caused by the substitution of conserved residues within alpha helix 1 of the DNA-binding domain (W95R and A104P), and dominant-negative mutations caused by a premature truncation which mimics the shortening caused by naturally occurring mutations (Q161*, E169*, G204* and C240*). Even though some interactions are lost in some mutant conditions, a large number of protein partners are retained. Different subsets of protein partners are retained depending upon the extent of SOX18 protein truncation. Mutations G204* and C240* are the closest counterparts to SOX18^RaOp^ as the DNA-binding HMG domain and homodimerization DIM domain is left intact, and the C-terminal TAD domain is disrupted. This indicates that SOX18^RaOp^ has the potential to directly compete for SOX18 homodimer formation and protein partner recruitment to not only block SOX18 transcriptional activity but those of its interactors as well. Further, the difference in the protein partner recruitment for different mutants would contribute to the variance in phenotype severity observed between mutants.

**Fig. 3.**
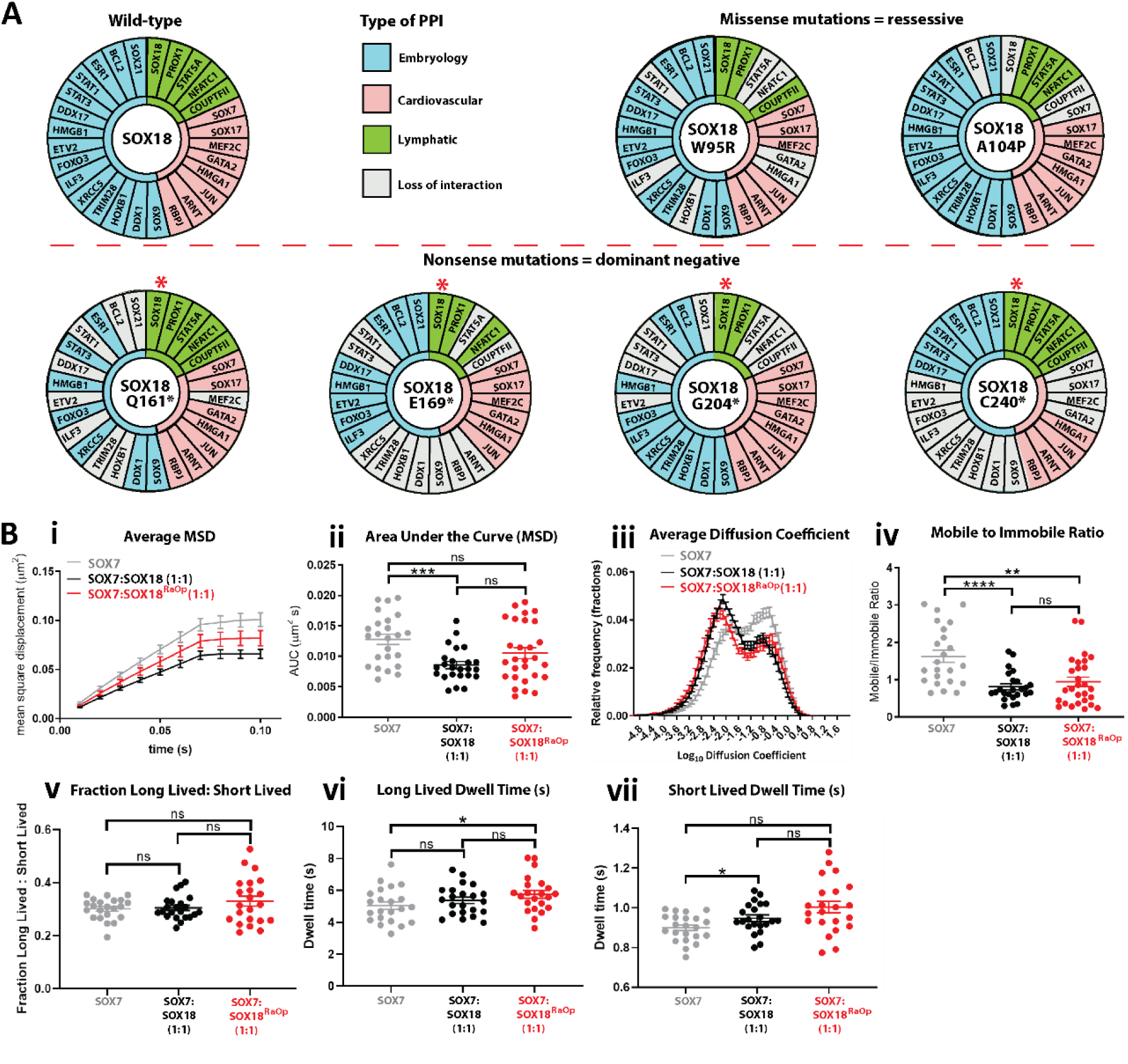
The SOX18^RaOp^ dominant-negative mutant recruits many SOX18 protein partners to form nonfunctional complexes. (**A**) (**i**) AlphaScreen assay to assess pair-wise protein-protein interactions in presence of different SOX18 human mutations. Loss of interaction shown in grey. All dominant-negative mutants retain their ability to form a dimer with wild-type SOX18 (red asterisks). (**B**) Quantification of the dynamics of HALO-SOX7 (grey), HALO-SOX7 with untagged-SOX18 (black) in a 1:1 ratio and HALO-SOX7 with untagged-SOX18^RaOp^ in a 1:1 ratio (red). (**i**) the average mean square displacement (MSD; μm^2^), (**ii**) the area under the curve for the average mean square displacement (μm^2^s), (**iii**) the average diffusion coefficient (μm^2^/s), and (**iv**) the mobile to immobile ratio. Threshold to classify mobile and immobile molecules is Log_10_D = -1.5. Values for the mean ± s.e.m. are shown. Average number of trajectories: HALO-SOX7 = 2737, HALO-SOX7:SOX18 (1:1) = 2370, HALO-SOX7:SOX18^RaOp^ = 2438. n = 24 for HALO-SOX7, n = 21 for HALO-SOX7:SOX18 (1:1), n = 27 for HALO-SOX7:SOX18^RaOp^ (1:1) (N = 3). t-test (two-tailed, unpaired). ** P<0.01 *** P<0.001, ns = non-significant (P>0.05). (**v**) the fraction of long lived to short lived immobile events, duration of (**vi**) long-lived immobile and (**vii**) short-lived immobile events. Values for the mean ± s.e.m. are shown. n = 22 for HALO-SOX7, HALO-SOX7:SOX18 (1:1) and HALO-SOX7:SOX18^RaOp^ (1:1) (N = 3). Mann Whitney U-test (two-tailed, unpaired). * P<0.05, ns = non-significant (P>0.05).

The current hypothesis on the dominant-negative mode of action of the SOX18^RaOp^ protein is its ability not only to disrupt SOX18 wild-type protein activity but more broadly to interfere with closely related SOXF family members (SOX7 and SOX17), which in turn inhibits any redundancy mechanism. As shown in Fig. 3A, SOX7 and SOX17 are recruited by the majority of non-functional SOX18 mutants. This proposed molecular mechanism would explain why ragged mice exhibit severe vascular defects whereas the SOX18 knockout mice are devoid of cardiovascular defects (*15, 21, 45*) in certain genetic backgrounds. To analyze the level of interference of SOX18^RaOp^ on other SOX members, we performed SMT on HALO-SOX7 and HALO-SOX17 to quantify changes in their chromatin-binding dynamics in the presence of either untagged-SOX18 or untagged-SOX18^RaOp^ (Fig. 3B, and Fig. S1 to Fig. 3). Overall, the chromatin-binding behavior of SOX7 and SOX17 changes in the presence of SOX18. Interestingly, similar changes in chromatin binding dynamics are also observed upon the addition of SOX18^RaOp^. Compared to SOX7 by itself, upon the addition of SOX18 or SOX18^RaOp^, SOX7 has a significantly higher chromatin-bound fraction and longer specific dwell time (Fig. 3B). These changes are shared by SOX17, which in addition also has a significantly lower overall mobility (Fig. S1 to Fig. 3). This shows that SOX18^RaOp^ can in part compete with SOX18 for SOX7 and SOX17, resulting in non-functional SOX7/SOX18^RaOp^ and SOX17/SOX18^RaOp^ heteromers on the chromatin.

### The SOX18^RaOp^ mutation perturbs the oligomeric state of SOX18 and affects chromatin-binding dynamics of dimer-specific protein partners

One common PPI across all dominant-negative mutations is the wild-type SOX18 protein (Fig. 3A, asterisks), which is consistently recruited by its mutant counterpart. This suggests that in presence of the SOX18^RaOp^ protein a mixture of different homo- and hetero-dimers are coexisting (SOX18/SOX18, SOX18/SOX18^RaOp^, SOX18^RaOp^/SOX18 and SOX18^RaOp^/SOX18^RaOp^), with a bias towards non-functional protein complexes. Previous work has reported that a functional feature of the SOX18 protein is its ability to form homodimers – a molecular state tightly associated with an endothelial-specific transcriptional signature (*46*). A key characteristic of TF activity is its ability to modulate mRNA transcription rate by communicating with basal transcriptional machinery. In order to further validate the functional relevance of the homodimer complex, we compared the overlap of total SOX18 and SOX18 homodimer only ChIP-seq peaks with active or repressive histone marks, and RNA polymerase II binding regions (Fig. 4A and 4B) taking advantage of data sets from HUVECs generated by the ENCODE consortium (*47–50*). This showed an enrichment of the SOX18 ChIP-seq peaks which harbor a SOX dimer motif (Inverted repeat 5; IR 5 (*46*)) with a broad range of histone marks and RNA polymerase II. This observation indicates that the SOX18 homodimer localizes to transcriptionally active sites engaged in either activation or repression.

**Fig. 4.**
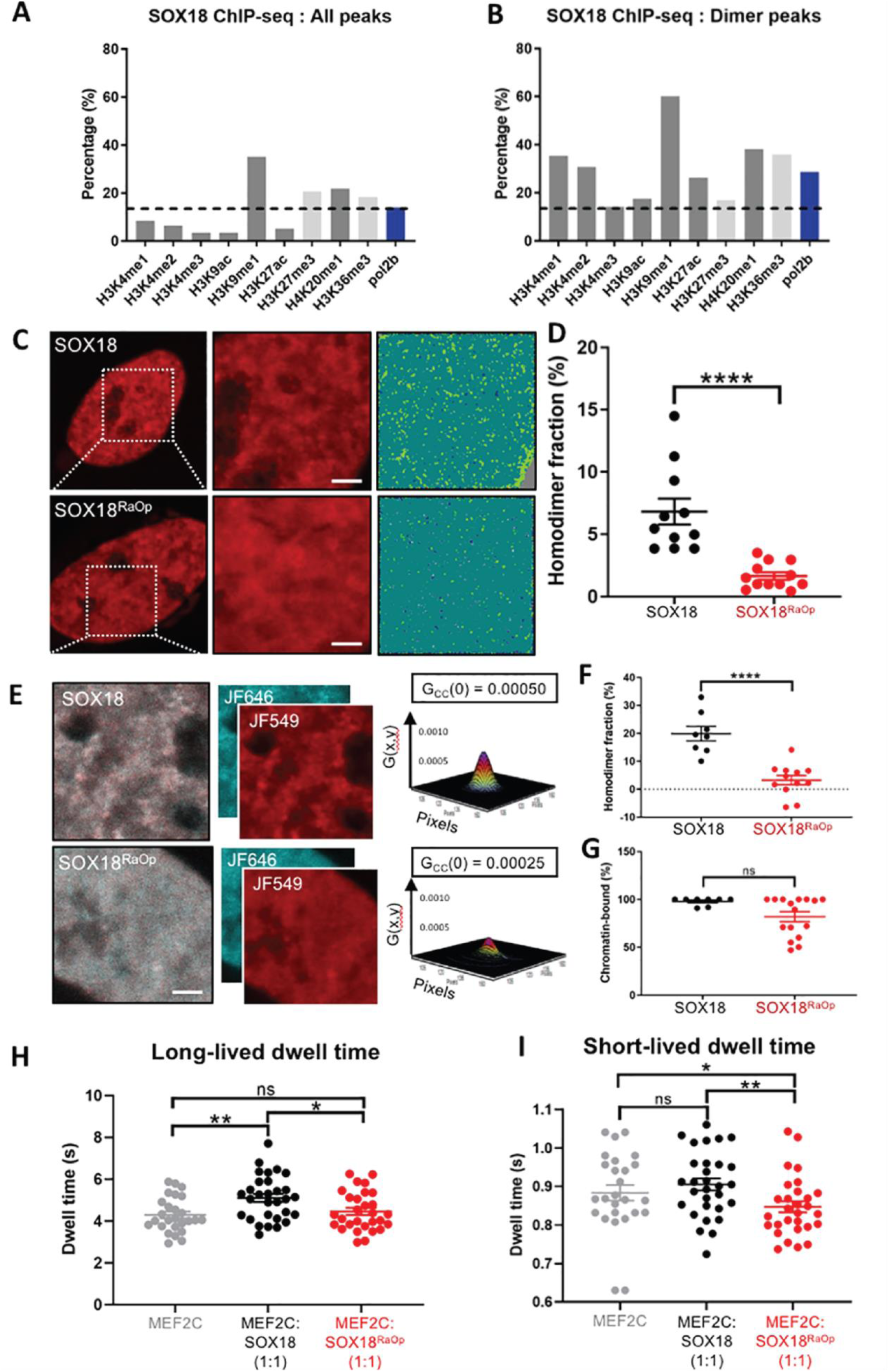
The SOX18^RaOp^ dominant-negative mutant has compromised homodimerization and impairs the scanning behavior of SOX18 dimer-specific protein partner MEF2C. (A) Overlay of SOX18 ChIP-seq peaks, or **(B)** SOX18 ChIP-seq peaks containing an IR5 dimer motif, with activating histone marks (dark grey), repressive marks (light grey) and RNA polymerase II (blue) in HUVECs. (**C**) Number and brightness (N&B) analysis. Nuclear regions in which N&B analysis was performed is outlined by a white box. Scale bar = 2 µM. Brightness maps indicate the oligomeric distribution of HALO-SOX18 and HALO-SOX18^RaOp^ with monomers/heterodimers (dark green), homodimers (light green), and an absence of molecules (dark blue). (**D**) Quantification of N&B analysis. HALO-SOX18 n = 11 and HALO-SOX18^RaOp^ n = 12 (N = 2). Values for the mean ± s.e.m. are shown. Mann Whitney U-test (two-tailed, unpaired). **** P<0.0001. (**E**) Cross-raster image correlation spectroscopy (cRICS) analysis of (**top row**) two-color (JF549 and JF646 dyes) HALO-tagged SOX18 and (**bottom row**) HALO-tagged SOX18^RaOp^. Both the JF549 and JF646 channels have been merged and cross-correlated to obtain 3-dimensional cross-RICS functions. Scale bar = 2 µM. (**F**) cRICS quantification to obtain the fraction of homodimers for HALO-SOX18 (black) and HALO-SOX18^RaOp^ (red). n = 8 and n = 12 (N = 3). Values for the mean ± s.e.m. are shown. Mann Whitney U-test (two-tailed, unpaired). **** P<0.0001. (**G**) cRICS quantification to obtain the fraction of homodimers that are bound to chromatin for HALO-SOX18 (black) and HALO-SOX18^RaOp^ (red). n = 8 and n = 16 (N = 2). Values for the mean ± s.e.m. are shown. Mann Whitney U-test for slow tracking (two-tailed, unpaired). ns = non-significant. (**H**) Single molecule tracking (SMT) to obtain the long-lived dwell times of HALO-MEF2C (grey), HALO-MEF2C with untagged SOX18 in a 1:1 ratio (black) and HALO-MEF2C with untagged SOX18^RaOp^ in a 1:1 ratio (red). Values for the mean ± s.e.m. are shown. n = 26 for HALO-MEF2C, n = 30 for HALO-MEF2C:SOX18 (1:1) and n = 29 for HALO-MEF2C:SOX18^RaOp^ (1:1) (N = 3). Mann Whitney U-test for slow tracking (two-tailed, unpaired). * P<0.05, ** P<0.01, ns = non-significant (P>0.05). (**I**) Slow tracking SMT to obtain the short-lived dwell times of HALO-MEF2C (grey), HALO-MEF2C with untagged SOX18 in a 1:1 ratio (black) and HALO-MEF2C with untagged SOX18^RaOp^ in a 1:1 ratio (red). Values for the mean ± s.e.m. are shown. n = 26 for HALO-MEF2C, n = 30 for HALO-MEF2C:SOX18 (1:1) and n = 29 for HALO-MEF2C:SOX18^RaOp^ (1:1) (N = 3). Mann Whitney U-test (two-tailed, unpaired). * P<0.05, ** P<0.01, ns = non-significant (P>0.05).

We next performed N&B and cross-RICS (cRICS) in order to observe the spatial distribution and chromatin-bound fraction of SOX18 homodimers. In order to label as many HALO-SOX18 and HALO-SOX18^RaOP^ molecules as possible whilst retaining fluctuations in fluorescence intensity necessary for these techniques, we chose cells with low expression levels, and used 1 µM of JF549 dye for N&B experiments, and 500 nM of JF549 and JF646 dyes for cRICS experiments as 1 µM total dye was shown to saturate HALO-SOX18 (Fig. S1 to Fig. 4). These dye concentrations are much higher than the one used for SMT experiments (2 nM), however unlike N&B and cRICS which uses confocal microscopy and can tolerate high levels of molecule detections, SMT uses oblique illumination and relies on sparse labelling in order to decrease background fluorescence and distinguish the trajectories of individual molecules without significant overlap. Here, we have calibrated the brightness of the monomeric fraction by using SOX7 (Fig. S2 to Fig. 4) as no SOX7 homodimers were detected previously for this TF by AlphaScreen (*46*) or indirectly via ChIP-seq analysis since no IR5 dimer motif is enriched in SOX7 ChIP-seq peaks (*42, 46*). This N&B approach revealed that even with low levels of homodimer detected at low HALO-SOX18 expression levels, SOX18 was found to form homodimers clusters (local enrichment of homodimers) throughout the nucleus (Fig. S2 to Fig. 4). This clustering was found to increase further at higher levels of SOX18 expression, with the first evidence of a higher-order oligomeric form (more than 3 molecules in a complex) for SOX18 within homodimer clusters (Fig. S2 to Fig. 4). By contrast, higher concentrations of SOX7 did not result in the formation of more homodimers as expected of a mono/heterodimeric protein (Fig. S2 to Fig. 4). N&B analysis comparing the oligomeric profile of HALO-SOX18 and HALO-SOX18^RaOp^ showed that ∼7 % of total SOX18 is SOX18 homodimers, whereas the homodimer population for SOX18^RaOp^ is significantly reduced at ∼2 % of total SOX18^RaOp^ (Fig. 4C and 4D).

To further validate the reduction of SOX18^RaOp^ homodimers observed in N&B, and to obtain the fraction of chromatin-bound homodimers we performed cRICS analysis on HALO-SOX18 using two spectrally distinct Halo-tag fluorophores (JF549 and JF646) (Fig. 4E-G). Colocalization and co-movement of JF549 and JF646 tagged HALO-SOX18 dimers was observed for cRICS, thus validating the presence of the dimer in the N&B and cRICS assays, and quantification of this confirmed that ∼20 % of SOX18 molecules are homodimers, and identified that the majority if not all SOX18 homodimers are bound to the chromatin (∼98 %) (Fig. 4D). The detection of a higher percentage of SOX18 homodimer than what was reported for N&B, is due to the higher sensitivity of cRICS. Strikingly, the fraction of HALO-SOX18^RaOp^ homodimer was significantly depleted to approximately 3 %, confirming that SOX18^RaOp^ forms less homodimers than SOX18. In support of the decreased HALO-SOX18^RaOp^ homodimer fraction observed for cRICS, a brightness aggregation assay performed using different temperatures showed that the human dominant-negative mutants most similar to SOX18^RaOp^ (G204* and C240*) required a higher temperature than SOX18 to form aggregates, and is therefore less likely to homodimerize (Fig. S3 to fig. 4). As it has been shown by AlphaScreen that SOX18 is recruited by all dominant-negative form of SOX18 mutants (Fig. 3A), this failure to form SOX18^RaOp^ homodimers likely skews the formation of non-functional SOX18/SOX18^RaOp^ complexes, thus perturbing wild-type SOX18 function even further. All together the N&B and cRICS analysis further confirm the impairment of SOX18^RaOP^ molecular behavior on a genome-wide scale with a significant change in oligomeric state formation.

The validation of SOX18 dimer behavior was further controlled for by the use of a SOX18 mutant lacking the homodimerization domain (*46*) (HALO-SOX18^DIM^; Fig. S4 to fig. 4). N&B confirmed that deletion of the DIM domain almost completely abolished SOX18 homodimer formation (Fig. S4A-D to Fig. 4). Further, similar observations were made using the alpha helix 1 SOX18^AH1^ DNA-binding mutant (Fig. S4A-D to Fig. 4). Collectively these results show that SOX18 dimers require DNA binding for their formation or maintenance, therefore suggesting that dimerization is primarily mediated via a cooperative mechanism. Analysis at a higher resolution performed using the cRICS approach further validated these observations (Fig. S4E-H to Fig. 4), where on average no homodimer was detected for the SOX18^DIM^ or SOX18^AH1^, indicating that the low level of residual homodimers observed by N&B were coincidental and likely to correspond to co-binding events whereby two molecules were juxtaposed on the DNA (Fig. S4H to Fig. 4). Further, cRICS data validated that the majority ∼ 96 % of SOX18 homodimers are chromatin-bound (Fig. S4I to Fig. 4). Validation using SMT analysis confirmed that removing the homodimerization domain resulted in an increase in SOX18 mobility, and a decrease in the immobilized fraction (Fig. S4J to Fig. 4). Focusing on the immobile population of SOX18^DIM^ mutant, SMT showed that there were less long-lived immobilization events than short-lived, and that both immobilization events occurred for a shorter total period of time. This observation suggests a change in the genome scanning behavior of the SOX18 protein when its ability to form a dimer is compromised.

Previously we have reported that MEF2C is preferentially recruited by a SOX18 homodimer (*46*). This SOX18 protein partner is essential for vascular development (*51*). Here we set out to assess the role of SOX18 in this complex, and whether SOX18^RaOp^ directly alters the molecular kinetics of MEF2C (Fig. 4C), beyond the disruption of the chromatin-binding dynamics of the SOXF group. Measuring MEF2C behavior by SMT in the presence of SOX18 revealed an increase in the long-lived dwell times (Fig. 4C) but not the chromatin-binding fraction or long-lived binding fraction (Fig. S5 to Fig. 4), validating the recruitment of MEF2C and indicating that the SOX18 homodimer plays a role in stabilizing MEF2C on target sites. In comparison to this, SOX18^RaOp^ did not increase the long-lived dwell time of MEF2C, and rather decreased its short-lived dwell time, indicating that SOX18^RaOp^ fails to stabilize MEF2C on target sites, and disrupts its search for target sites along non-specific chromatin.

This study establishes a set of molecular rules which are necessary to drive genome-wide perturbation in the context of transcription factor dominant-negative mutation and therefore instruct the phenotypic outcome of a genetic disease. Hallmarks of the dominant-negative mechanism are characterized by the capacity to disable key features of SOX18 activity: 1) via interference with chromatin sampling behavior thought increased dwell times of non-functional complexes at target sites, 2) compromised oligomerization, and 3) broadly poisoning the SOX18 interactome which in turn impacts other TF networks.

## Discussion

In this study, we uncovered several core biophysical properties that define how the key vascular and hair follicle regulator SOX18 navigates the genome. Further, we describe the multifaceted way by which a dominant-negative SOX18 mutant interferes with its wild-type counterpart to perturb this search pattern on the chromatin while poisoning other TF networks via protein-protein interactions (Fig. 5). Our findings explain at the molecular level on a genome-wide scale the etiology of a rare disease which is underpinned by a non-functional TF.

**Fig. 5.**
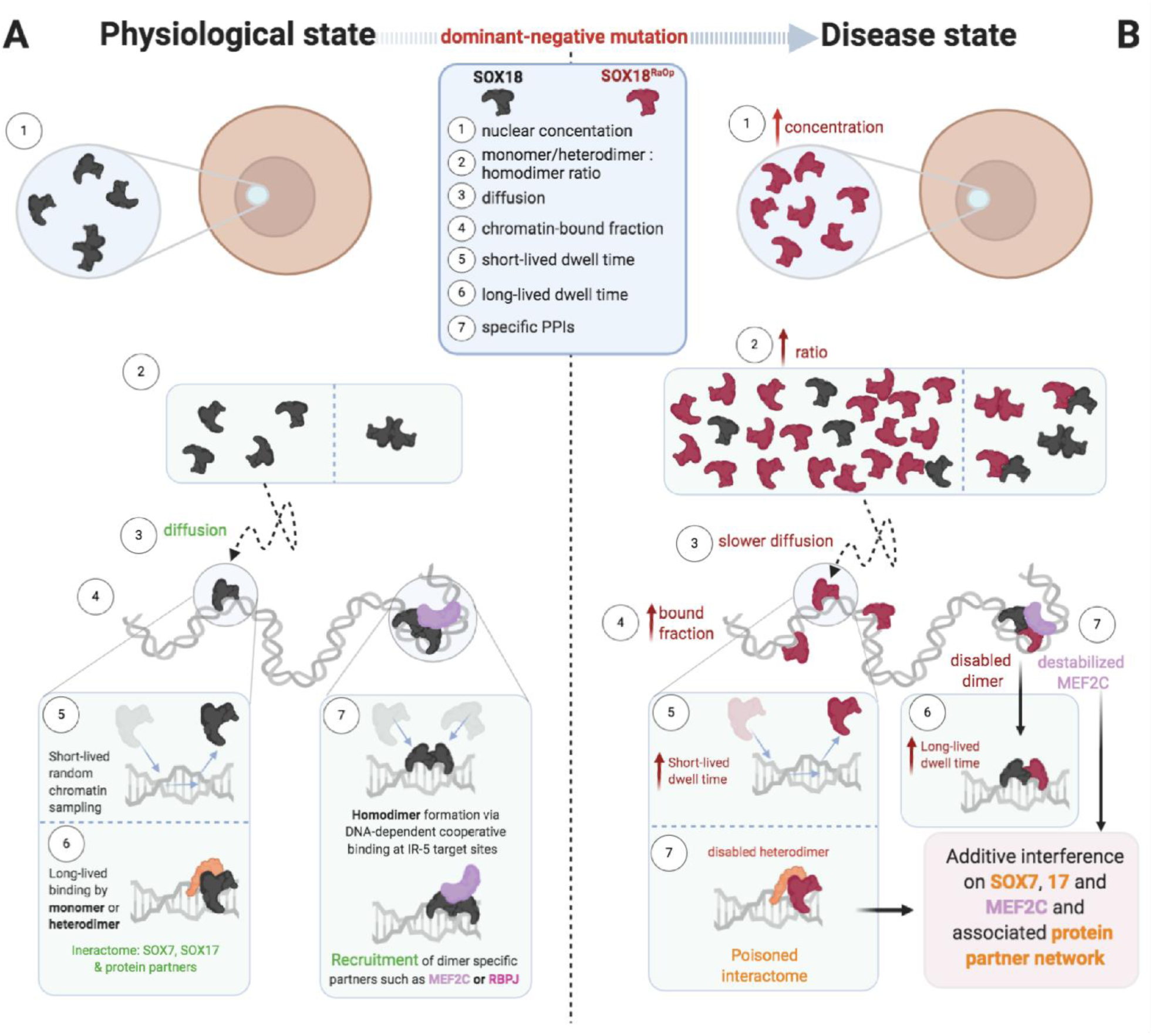
Components that make up the behavior of SOX18 in a physiological state, and how these regulatory layers are impacted by a dominant-negative SOX18 mutant to cause potent transcriptional repression. Created with BioRender.com (**A**) SOX18 transcription factor (black) behavior in a normal physiological state. (**1**) SOX18 is present at physiological levels within the nucleus (**2**) and exists in different oligomeric states. (**3**) SOX18 exhibits diffusion throughout the nucleus in between immobile events on the chromatin. (**4**) The SOX18 chromatin-bound fraction represents around 20 % of its population. (**5**) SOX18 dwells for a short period of time (less than 1 second) on non-specific sites as it samples the chromatin to identify target sites. **6)** SOX18 binds to target sites where it remains for longer (a few seconds). **7)** SOX18 homodimer formation and maintenance are DNA-dependent via cooperative binding to recruit homodimer-specific partners such as MEF2C or RPBJ. (**B**) SOX18^RaOp^ (red) is a dominant-negative mutant of SOX18 and exhibits aberrant behaviors leading to large scale transcriptional dysregulation. The interference of SOX protein partners by SOX18^RaOp^ is additive hence disrupting multiple regulatory networks.

The chromatin-binding stability of a TF is dictated by the protein-protein, protein-DNA and protein-RNA interactions that it forms (*1, 2*). In support of this DNA-binding and homodimerization SOX18 mutants (SOX18^DIM^ and SOX18^AH1^) showed a significant reduction in their chromatin-binding capabilities. Surprisingly, the opposite effect was observed for SOX18^RaOp^, indicating that the chromatin-binding behavior of this mutant is likely due to an increase in chromatin-binding stability, conferred to the wild-type protein upon interaction. In further support of this, despite being expressed at comparable levels with SOX18 throughout the entire cell, more SOX18^RaOp^ was retained within the nucleus. This increased stability may be explained by the change in the iso-electric point of SOX18^RaOp^ (i.e. 10.4) which is much higher than its wild-type counterpart (i.e. 7.3) (*52*). The DNA-binding HMG domain is already extremely positively-charged in order to bind negatively-charged DNA efficiently, so here we proposed that overall SOX18^RaOp^ acts as a chromatin “magnet”, which in turn disables a higher number of sites recognized as “specific” and its dwell times. The scrambling and premature truncation of the transactivation domain of SOX18^RaOp^ causes the overall charge to become skewed in favor of the positively charged DNA-binding HMG domain rich in high pKA amino acids, therefore likely increasing its affinity for negatively charged DNA.

Through some unknown mechanism, SOX18^RaOp^ interferes with the ability of other SOXF members (SOX7 and SOX17) to rescue the phenotype. We hypothesized that this interference may be reflected by changes in their chromatin-binding dynamics in the presence of SOX18^RaOp^ as compared to SOX18. We observed that SOX18^RaOp^ altered the chromatin-binding behavior for both SOXF factors in a similar way to SOX18, suggesting that SOX18^RaOp^ has the ability to compete with SOX18 for SOXF factors. The recruitment of SOXF factors by SOX18^RaOp^ to form a non-functional complex is supported by PPI data showing that SOXF factors are recruited by SOX18 and the majority of SOX18 mutants assessed here.

To investigate a potential interference with a shared protein partner, we assessed the ability of SOX18 and SOX18^RaOp^ to alter the chromatin-binding dynamics of MEF2C. Rather unexpectedly, SOX18 increased MEF2C stability on target sites, whereas SOX18^RaOp^ destabilized MEF2C on both specific target sites, and during its search along non-specific chromatin sites. MEF2C has previously been reported to be preferentially recruited by a SOX18 homodimer (*46*), and, since SOX18^RaOp^ has a significant reduction in chromatin-bound homodimers, this suggests that it would be unable to efficiently stabilize MEF2C on active chromatin sites. Importantly, this molecular scenario is not at play in the context of SOX18 targeted gene disruption whereby a lack of SOX18 does not interfere directly with the search pattern mechanism of MEF2C, hence limiting its range of interference.

It remains unclear as to why some interactions are strengthened and others weakened, although given that different PPIs involve different protein subdomains this is not entirely unexpected. The mutant protein retains multiple protein partners, suggesting that the observed negative effect on transcription could potentially be amplified via interference with multiple regulatory hubs. This hypothesis is supported by the observation that the PPI profile of the different Ragged mutations correlates with disease phenotype severity. The longer the mutant protein, the more severe the phenotype, in other words as the mutant protein gets shorter it loses the ability to poison other protein complexes.

Although we have made considerable progress towards uncovering the molecular mechanisms that underpin a genetic disorder, it remains unknown as to how these changes directly relate to the mutant phenotypes observed in humans and mice. Genetic studies have shown that SOX18 directly controls the transcription of other key genetic pathways, such as Notch1 for arterial specification (*26, 27*), or Prox1 for lymphangiogenesis (*16*). Based on this, the question posed could be answered by directly measuring at a single locus (at a particular enhancer or promoter of Notch1 or Prox1 for instance) the chromatin-binding dynamics of SOX18 verses SOX18^RaOp^. At present, technological limitations relating to the sparse labelling approach required to perform SMT do not allow for this type of resolution. The number of events at a single locus is simply too low and not compatible with a fast acquisition rate (10 ms).

Prior to this study, only mathematical models to describe the broad spectrum interference of a dominant-negative TF have been developed (*5*). These models used two main variables: the level of allelic expression combined with the genetic configuration (homozygous or heterozygous). Here we reveal seven quantifiable components: nuclear concentration, the ratio of monomers/heterodimers to homodimers, the rate of protein diffusion, the chromatin-bound fraction, dwell times at specific target sites and non-specific sites, and protein partner recruitment. Further to this, we show that not only does the dominant-negative protein interfere with its wild-type counterpart, but it also interferes with the behavior of other classes of TF such as MEF protein. This extends the interference mechanism beyond just perturbing its own regulatory network, since the mutant affects the regulatory hubs belonging to its protein partners. This level of interference was not appreciated before and even less so demonstrated at the experimental level.

The question arises as to how applicable our model is other TF families. Here we propose two criteria to predict whether mutations would cause any TF to broadly interfere with transcription in SOX18^RaOp^-like fashion. Firstly, the dominant-negative mutation should leave the DNA-binding domain intact and disrupt the transactivation domain to remove its transcriptional capability whilst still enabling key protein partner recruitment. Secondly, the wild-type form should form DNA-dependent homodimers. Based on these assumptions, we believe that this model will not only apply to other SOX members with dominant-negative TF counterparts, but also to TFs of other families that satisfy these criteria.

In conclusion, by combining imaging techniques with genomics and proteomics assays, we were able to quantitate the effects of a dominant-negative TF on a genome-wide scale. We demonstrate that this broad interference is likely mediated by multiple parameters that directly relate to wild-type protein activity but also expand to other regulatory hubs engaged via PPIs. Looking ahead with the advent of new ways to visualize specific genomic locations or measure transcription rate in real time, the next step will be to correlate changes in TF activity measured in real time with their corresponding transcriptional output. This type of approach combined with multi-omics analysis will better our understanding of the molecular basis of gene regulation.

## Materials and Methods

### Plasmids

A list of plasmids used in this study is provided in Table S1.

pReceiver-M49(HaloTag-SOX18WT) was obtained from GeneCopoeia, and subsequently used as a template to generate all other HaloTag constructs using the In-Fusion HD Cloning Kit (Takara Bio USA, Inc), with the exception of pReciever-M49(HaloTag-SOX17) which was also obtained from GeneCopoeia. Protein expression for all plasmids is driven by the CMV-enhancer promoter.

Alpha-helix 1 of the SOX HMG domain was removed from pReceiver-M49(HaloTag-SOX18WT) using a combination of circular polymerase extension cloning (CPEC) with In-Fusion cloning to generate the pReceiver-M49(HaloTag-SOX18WTΔAH1) construct.

The homodimerization domain consisting of 50 amino acids (155-199) adjacent to the C-terminal NLS was removed from pReciever-M49(HaloTag-SOX18WT) using a combination of circular polymerase extension cloning (CPEC) with In-Fusion cloning to generate the pReceiver-M49(HaloTag-SOX18WTΔDIM) construct.

### Western blotting

Western blotting was used to assess the endogenous or overexpressed SOX18 protein level in HeLa cells and HUVECs. Cells were seeded, and either transfected (using 1 µg of expression plasmid to 3 µl of X-tremeGENE 9) or left untransfected, and harvested for either whole cell lysates or nuclear extracts before subjecting to SDS-PAGE and Western Blotting with a human anti-SOX18 antibody (sc166025 from Santa Cruz), anti-HaloTag antibody (G9281 from Promega), or housekeeping control anti-β-actin antibody (A5441 from Sigma).

### Luciferase assay

7000 Monkey kidney fibroblast-like COS-7cells were seeded per well in gelatin-coated 96- well plates (Gibco DMEM, Cat# 11995073, 10 % v/v heat-inactivated foetal bovine serum, 1 % L-glutamine, penicillin, streptomycin). Cells were maintained at 37 °C, in a 5 % CO_2_ controlled atmosphere. After 24 hours, a four-hour transfection with murine plasmids, pGL2-Basic (Promega) Vcam-1 promoter construct (VC1889; 40ng per well), with and without HALO-SOX18 and/or untagged-SOX18^RaOp^ was performed. (HALO-SOX18 alone = 30 ng/well, SOX18^RaOp^ alone = 10 ng/well, 30:1 ratio = 30 ng HALO-SOX18: 1 ng SOX18^RaOp^ etc., 40ng of each plasmid per well in 10 µL of premix X-tremeGENE HP DNA transfection reagent, Roche/Sigma, Cat# 06 366 236 001, 1:4 DNA:Xtreme ratio) (*43*).

### Cell culture

HeLa cells were a gift from Professor Geoffrey Faulkner (Queensland Brain Institute/Translational Research Institute, St Lucia, Queensland, Australia). Cells were cultured in Dulbecco’s Modified Eagle Media (DMEM, Glibco) supplemented with 10 % Fetal Bovine Serum (FBS, GE Healthcare), 1 % GlutaMAX (Glibco) and 1 % MEM Non-Essential Amino Acids (MEM NEAA, Glibco). Cells were maintained at 37 °C with 5 % CO_2_.

### Cell seeding and transfection

HeLa cells were seeded at a density of 155,000 cells/ml in 35 mm glass coverslip dishes (P35G-1.5-20-C, Matek) coated with 1 % gelatin 24 h prior to transfection. Transfections were performed using the X-tremeGENE 9 Transfection Reagent kit (Roche) to introduce 1-2 μg of plasmid DNA as per manufacturer’s instructions, using FluoroBrite DMEM (Glibco) supplemented with 1 % GlutaMAX (Glibco) as the low serum transfection media. Cells were incubated at 37 °C with 5 % CO_2_ for 24 h prior to imaging.

### Single Molecule Tracking – Imaging

Immediately prior to imaging cells were washed twice and replaced with Fluorobrite DMEM (Glibco) imaging media. JF549 was a gift form Dr. Luke Lavis (Janelia Research Campus, Howard Hughes Medical Institute, Ashburn, Virginia, United States). 2 drops/mL of NucBlue Live ReadyProbes Reagent (Hoechst 33342) was added directly to the media and cells were incubated for 5 min at 37 °C with 5 % CO_2_, prior to adding 2 nM of JF549 Halo-tag dye directly to the media and cells and incubation for a further 15 min at 37 °C with 5 % of CO^2^. Following incubation, cells were washed twice and replaced with Fluorobrite DMEM (Glibco).

Images were acquired on an Elyra single molecule imaging (PALM/STORM) total internal reflection fluorescence (TIRF) microscope, with an Andor 897 EMCCD camera, SR Cube 05 RL – BP 420-480 / BP570-640 / LP 740 filter set and 100 X oil 1.46 NA TIRF objective using ZEISS ZEN blue software.

Cells were imaged using a 561 nm excitation laser at ∼11 µW to perform two different acquisition techniques; fast tracking which uses a 20 ms acquisition speed to acquire 6000 frames without intervals, and slow tracking which uses a 500 ms acquisition speed to acquire 500 frames without intervals.

### Single molecule tracking – Fast tracking (20ms) analysis

Fast tracking (20ms) SMT raw image stacks were cropped in ImageJ prior to analysis to reduce their file size and therefore minimize analysis time. Fast tracking SMT data was then analyzed using the PalmTracer plugin developed for Metamorph by the Sibarita group at the University of Bordeaux (*53*). As described by Bademosi and colleagues (*54*), here we used PALM-Tracer to localize and track molecules in order to obtain their trajectories, and to calculate the mean square displacement (MSD) and diffusion coefficient (D) for each trajectory. For molecule localization, we used a watershed of size 6. To reduce non-specific background, trajectories were filtered based on a minimum length of 8 and a maximum length of 1000, and a maximum travel distance of 5 µM. For visualization, a zoom of 8 with a fixed intensity and size of 1 was used. A spatial calibration of 100 nm and a temporal calibration of 20 ms was used. MSD fitting was performed using a log scale with a length of 4 and a step number of 1.

Color-coding of fluorescence intensity, diffusion coefficient and trajectory heatmaps was performed using ImageJ. Fluorescence intensity heatmaps range from black (no molecules detected) to white (highest density detected). Diffusion coefficient heatmaps range from white showing the highest diffusion coefficient to black showing the lowest diffusion coefficient. Trajectories are arbitrarily colored to help discriminate them visually.

Analysis files produced by PALM-Tracer were used as input for AutoAnalysis_SPT software (https://github.com/QBI-Software/AutoAnalysis_SPT/wiki) developed for the Meunier Laboratory at the Queensland Brain Institute by Dr Liz Cooper-Williams. AutoAnalysis_SPT software compiles the results obtained for each cell to obtain the average MSD, calculates the average area under the curve of the MSD for each cell, generates a histogram showing the distribution of the different diffusion coefficients, and calculates the mobile to immobile ratio for each cell. Here we used 10 MSD points, with a time interval of 0.02 s (20ms acquisition time) and included trajectories with a minimum diffusion coefficient of -5, and a maximum diffusion coefficient of 1. For the diffusion coefficient histogram, we chose a bin width of 0.1, and a mobile to immobile threshold of -1.5, determined mathematically using the equation

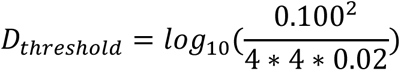

where 0.100 is the pixel size in nm, 0.02 is the acquisition time in seconds and 4 refers to the number of MSD points used for fitting.

Individual MSDs for mobile and immobile fractions were calculated manually by segregating MSDs for each cell based on their diffusion coefficient (less than -1.5 = immobile, greater than -1.5 = mobile), with trajectories with diffusion coefficients higher than 1 and lower than -5 excluded. In Graph Pad Prism (version 8.0) the AUC was calculated for each cell using default settings.

Values for the mean ± s.e.m. were plotted into Graph Pad Prism, and unpaired two-tailed t-tests were used to assess significance of the AUC for the MSD and the mobile to immobile ratio of the diffusion coefficient histogram. Cells with significant drift or less than 1000 trajectories were excluded from analysis, with the exception of HALO-SOX18^DIM^ for which cells with less than 800 trajectories were excluded. A robust regression and outlier removal (ROUT) outlier test using default settings (Q = 1 %) in Graph Pad Prism was performed to identify and remove outliers.

### Single molecule tracking – slow tracking (500ms) analysis

Slow tracking (500ms) SMT raw image stacks were cropped in ImageJ prior to analysis to reduce their file size and therefore minimize analysis time.

Slow tracking SMT data was analysed using a Matlab pipeline using Matlab version R2015a as previously published by Chen and colleagues (*2*). Here we used this Matlab pipeline to assess whether the immobile fraction consists of one or two types of dwell times (long-lived, a few seconds and short-lived, less than 1 s), calculate the fraction of long-lived to short lived dwell times and the length of time for which they occurred. This pipeline first uses the Matlab script SLIMfast.m to localise molecules, which is a modified version of the multiple trace tracking (MTT) algorithm reported by Sergé and colleagues (*55*). SLIMfast.m is available on the eLife website. SLIMfast was performed using an error rate of 10^-7^, a detection box of 7 pixels, a maximum number of iterations of 50, a termination tolerance of 10^-2^, a maximum position refinement of 1.5 pixels, an N.A. of 1.46, a PSF scaling factor of 1.35, 20.2 counts per photon, an emission 590 nm, a lag time of 500 ms and a pixel size 100 nm. Following this, the Matlab script Calculatelength_2fitting_v3.m (available on request) is used to calculate the lifetime of molecules for each cell, and fits one and two-component exponential decay curves to this data and using the equation

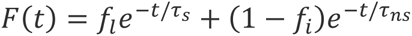

for two-component model derives the average dwell time for specific and non-specific fractions and the ratio between these. The values extracted from this were plotted into Graph Pad Prism with the mean ± s.e.m. are shown. Statistical significance was assessed using a two-tailed unpaired Mann Whitney U-test for slow tracking. Cells with significant drift were excluded from analysis. A robust regression and outlier removal (ROUT) outlier test using default settings (Q = 1 %) in Graph Pad Prism was performed to identify and remove outliers.

### ChIP-MS

Protein partners identified for SOX18 via ChIP-MS have been reported previously (*56*). These were used here as a basis for screening interactions with dominant-negative SOX18 mutants via AlphaScreen analysis.

### AlphaScreen

AlphaScreen assays were performed as previously described (*46*).

### GTEx analysis

Genes previously reported by ChIP-seq to contain regulatory elements with a SOX18 homodimer binding motif (IR-5) (*46*), and genes reported to contain a MEF2C binding motif (DMSO control) (*57*) were input into GTEx (*58*) online to obtain an expression matrix in tissues selected for their association with blood vessel and hair follicle development.

### Fluorescence fluctuation spectroscopy aggregation assay

Fluorescence fluctuation spectroscopy aggregation assays were performed as previously described (*59*).

### Confocal microscopy for fluorescence fluctuation spectroscopy

Confocal microscopy for fluorescence fluctuation spectroscopy was performed as previously described. All microscopy measurements were performed on an Olympus FV3000 laser scanning microscope coupled to an ISS A320 Fast FLIM box for fluorescence fluctuation data acquisition. A 60 X water immersion objective 1.2 NA was used for all experiments and live HeLa cells were imaged at 37 degrees in 5 % CO2. For single color fluorescence fluctuation spectroscopy experiments SOX7 (Fig. S2A and S2E to Fig. 4), SOX18 (Fig. 4C and 4D, and Fig. S2A-D and S2F to Fig. 4) SOX18 mutants (Fig. S2A-D to Fig. 4) and SOX18^RaOp^ (Fig. 4C and 4D) were labelled 15 minutes prior to imaging via direct addition of 1 µM of JF549 Halo-tag dye, where JF549 was excited by a solid-state laser diode operating at 561 nm. The fluorescence signal was then directed through a 405/488/561 dichroic mirror to remove laser light and the JF549 emission collected through a 550 nm long pass filter by an external photomultiplier detector (H7422P-40 of Hamamatsu) fitted with a 620/50 nm bandwidth filter. A 100-frame scan acquisition of the J549 signal was then collected by selecting a region of interest within a HeLa cell nucleus at zoom 20, which for a 256 x 256-pixel frame size resulted in a pixel size of 41 nm. The pixel dwell time was set to 12.5 µs, which resulted in a line time of 4.313 ms and a frame time of 1.108 s.

For the two color experiments (Fig. 4E-G, Fig. S2B-D to Fig.4, and Fig. S3E-H to Fig.4) SOX18, SOX7 and SOX18 mutants were co-labelled with 500 nM of JF549 as well as 500 nM of JF647 and these two dyes were excited by solid-state laser diodes operating at 561 nm and 640 nm (respectively). The fluorescence signal was then directed through a 405/488/561/640 dichroic mirror to remove laser light and the JF549 versus JF647 emission detected by two internal GaAsp photomultiplier detectors set to between the following bandwidths: JF549 570-620 nm, JF47 650-750 nm. A two channel 100-frame scan acquisition of the J549 and JF647 signal was then collected simultaneously employing the same acquisition settings described above for the single channel experiment.

### Number and brightness

Number and brightness (NB) analysis of frame scan acquisitions, was performed using a moment-based analysis described in previously published papers. Briefly, in each pixel of the frame scan we have an intensity fluctuation that has an average intensity (first moment) and a variance (second moment). The ratio of these two properties describes the apparent brightness (B) of the molecules that give rise to the intensity fluctuation. In the case of a photon counting detector, the true molecular brightness (*ɛ*) of the molecules are related to the measured apparent brightness (*B*) by

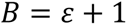

where 1 is the brightness contribution of the detector given that the photon-counting detector variance should be equal to the average intensity of the detector noise. Calibration of the monomeric brightness of JF549 enabled extrapolation of the expected brightness of different sized HALO-tagged protein oligomers and quantitation of the fraction of pixels within a given frame scan acquisition containing HALO-tagged protein monomers, dimers and oligomers. Artefacts due to cell movement or cell bleaching were subtracted from acquired intensity fluctuations via use of a moving average algorithm. All brightness calculations were carried out from the NB page in SimFCS from the Laboratory for Fluorescence Dynamics (www.lfd.uci.edu).

### Cross-correlation raster image correlation spectroscopy

Cells were imaged on an Olympus FV3000 microscope fitted with a heated CO_2_ stage at 37°C and 5 % CO_2_. EGFP fluorescence was excited with a 488 nm laser (Coherent Obis) at 0.1-0.4 % laser power. Emitted fluorescence was routed via the fiber port to a photon counting detector (ISS A320 FastFlim). Transfection yielded cells with a variety of expression levels (Fig. S2 to Fig. 4), and cells with a medium-low fluorescence intensity were selected for cRICS. To acquire cRICS data, 100 confocal images were acquired as a stream acquisition on a cell nucleus (zoom 20, 256×256 pixels, pixel size 41 nm, pixel dwell time 12.5 μs). cRICS data was analyzed in SimFCS subtracting a moving average of 10 frames. cRICS plots were fit to a 2-component 3D diffusion model in SimFCS fixing the pixel size (41 nm), line time (4.313 ms) and jumping 3 points in x. The beam waist, Wo = 0.26 μm was calibrated by measuring recombinant GFP in solution (D ∼ 90 μm^2^/s). The approximate number of molecules in the observation volume was calculated as

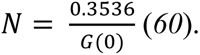

The average N for data in Fig. 4 and Fig. S2 to Fig. 4 is 403.5 molecules, which corresponds to a concentration of 44.7 nM given a confocal observation volume of 15 fL. Statistical analysis and plotting was performed using Graphpad Prism 8.0 software.

### Statistical Analysis

All statistical analyses were performed in Graph Pad Prism (version 8.0). A robust regression and outlier removal (ROUT) outlier test using default settings (Q = 1 %) in Graph Pad Prism was performed to identify and remove outliers. Significance was assessed for fast tracking SMT, N&B and cRICS analyses using unpaired two-tailed t-tests, and for slow tracking SMT analyses using two-tailed unpaired Mann Whitney U-tests. * P<0.05, ** P<0.01, **** P<0.0001, ns = non-significant (P>0.05).

## Acknowledgments

**General:** HeLa cells were a gift from Professor Geoffrey Faulkner (Translational Research Institute, Woolongabba, QLD, Australia). Halo-tag dyes JF549 and JF646 were a gift from Dr. Luke Lavis (Janelia Research Campus, Howard Hughes Medical Institute, Ashburn, VA, United States). PB53x EF1-Dendra2-RPB1Amr was a gift from Timothée Lionnet (Addgene plasmid # 81228 ; http://n2t.net/addgene:81228 ; RRID:Addgene_81228).

## Author contributions

M.F. and F.F. generated the original idea. M.F, A.M and E.H contributed intellectually around experimental design and data analysis. A.M performed and analysed the single molecule tracking experiments. J.L. and E.H. performed and analysed the number and brightness and raster image correlation spectroscopy experiments. W.L. performed the Western blot analyses. M.F. performed the GTeX analysis on SOX18 ChIP-seq data set. M.M., E.S. and Y.G. performed and analysed the AlphaScreen and fluorescence fluctuation spectroscopy aggregation assays. F.F. performed the luciferase assay. F.M. provided access to a microscope at QBI for single molecule tracking and a computer with software for data analysis. Z.L. provided access to cell culture, single molecule tracking training and facilities at HHMI, and access to in-house single molecule tracking Matlab code. H.L. provided cell culture, single molecule tracking and data analysis training. A.B. provided single molecule tracking training and performed some of the single molecule tracking experiments. F.F. provided cell culture training and assisted with the development of the SOX18 homodimerisation and Ragged dominant negative mode of action models. A.M. and P.K. wrote the manuscript which M.F., E.H., Z.L., F.M., and A.B. critically revised. M.F and E.H supported the work with funding. MF coordinated the study across the multiple laboratories.

## Data and materials availability

All raw data will be deposited on data dryad and made publicly available. The reagents used for the paper such as expression vectors or cell lines are available but subject to MTA with the University of Sydney or University of Melbourne.

## Supplementary Materials

### Supplementary Figures

**Fig. S1 to Fig. 1.**
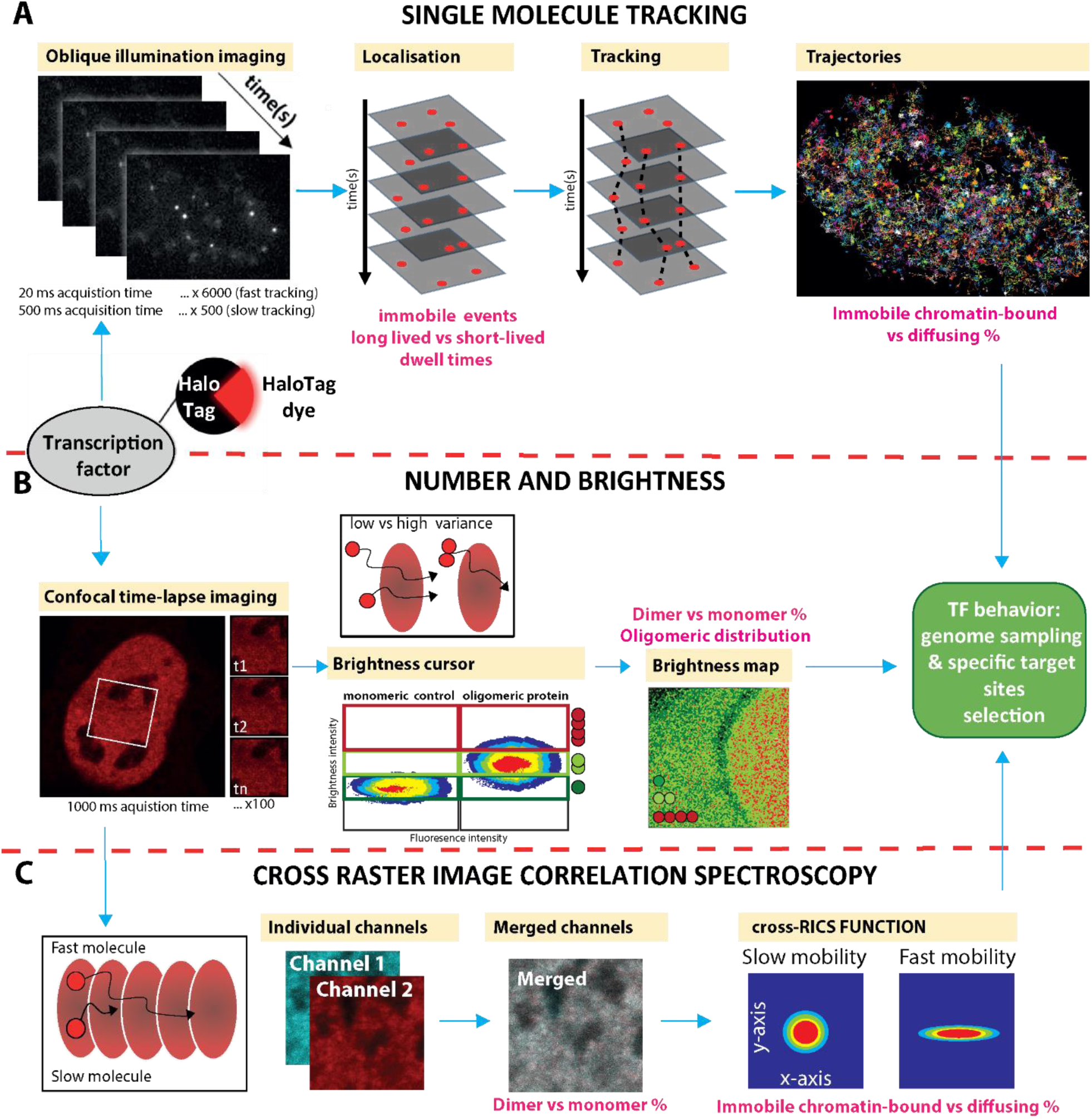
**Combination of single molecule-based assays in live cells to obtain the chromatin-binding dynamics, oligomeric distribution and chromatin-bound homodimer fraction of a transcription factor in order to uncover core components of its behavior.** Halo-tag labelling technology was used for all experiments. (A) **Single molecule tracking (SMT).** A series of images is taken using oblique illumination in order to penetrate the nucleus with a thin beam, greatly reducing background fluorescence and obtaining single molecule resolution. **Fast tracking SMT (20 ms acquisition times, 6000 frames):** in order to obtain the trajectories of each molecule, an acquisition time of 20 ms is used to capture diffusing molecules with faster mobilities in addition to immobile molecules. The location of each of the molecules in each of the frames is located (localization) and connected between frames (tracking) to obtain the trajectories of each molecule. Quantification of the trajectories gives immobile chromatin-bound and mobile diffusing fractions. **Slow tracking SMT (500 ms acquisition times, 500 frames):** within the immobile fraction of molecules, subpopulations of molecules with different dwell times exist. An acquisition time of 500 ms is used to decrease the signal of mobile diffusing molecules and enhance visualization of immobile chromatin-bound molecules to enable quantification of their dwell times and percentages. (**B**) **Number and brightness (N&B):** A region of interest is selected (white box outline), and a raster scan is performed using an acquisition time of 1000 ms in order to generate a confocal time series consisting of 100 frames. This acquisition time focuses analysis on the behavior of molecules with lower mobilities. For each pixel in this region of interest, the change in fluorescence intensity over time is measured and converted to fluctuations in molecular brightness, which are indicative of the average oligomeric state of molecules present in that pixel. A homodimer is twice as bright as a monomer or heterodimer (which cannot be distinguished here due to a single tag being used) and a higher-order oligomer is brighter than the dimeric state. Brightness is calibrated using a transcription factor that does not form detectable homodimers. This is used as a reference placing a brightness cursor around the brightness and fluorescence intensity of the monomer population (dark green). A homodimer cursor (light green) of the same height is placed directly above this, and a higher order oligomer cursor (3 or more TFs in a complex; red) is placed above this. The largest oligomeric form present within the pixel is depicted using a color code to generate a brightness heat map which shows the distribution of the different oligomeric states of the TF throughout the cell. From this heat map it is possible to infer the ability for the TF to form homodimers and higher-order oligomers, and the ability of these to generate clusters (higher local concentration of homodimers and higher-order oligomers). Shown in this example are monomers in dark green, homodimers in light green and higher order oligomers in red. (**C**) **Cross-raster image correlation spectroscopy (cRICS).** A region of interest is selected (white box outline), and a raster scan is performed using an acquisition time of 1000 ms in order to generate a confocal time series consisting of 100 frames. This acquisition time focuses analysis on the behavior of molecules with lower mobilities. For each pixel, the fluorescence intensity is obtained and then spatially correlated to the fluorescence intensity of each pixel in the rest of that confocal frame. The average spatial correlation of all the frames is then obtained from which a RICS function can be obtained and quantified to give the mobility of the molecules and the fraction that are immobile and chromatin bound. cRICS analysis using two spectrally distinct Halo-tag dyes validates the presence of homodimers as they diffuse together and enables the percentage of homodimers and chromatin-bound fraction of homodimers to be extracted.

**Fig. S2 to Fig. 1.**
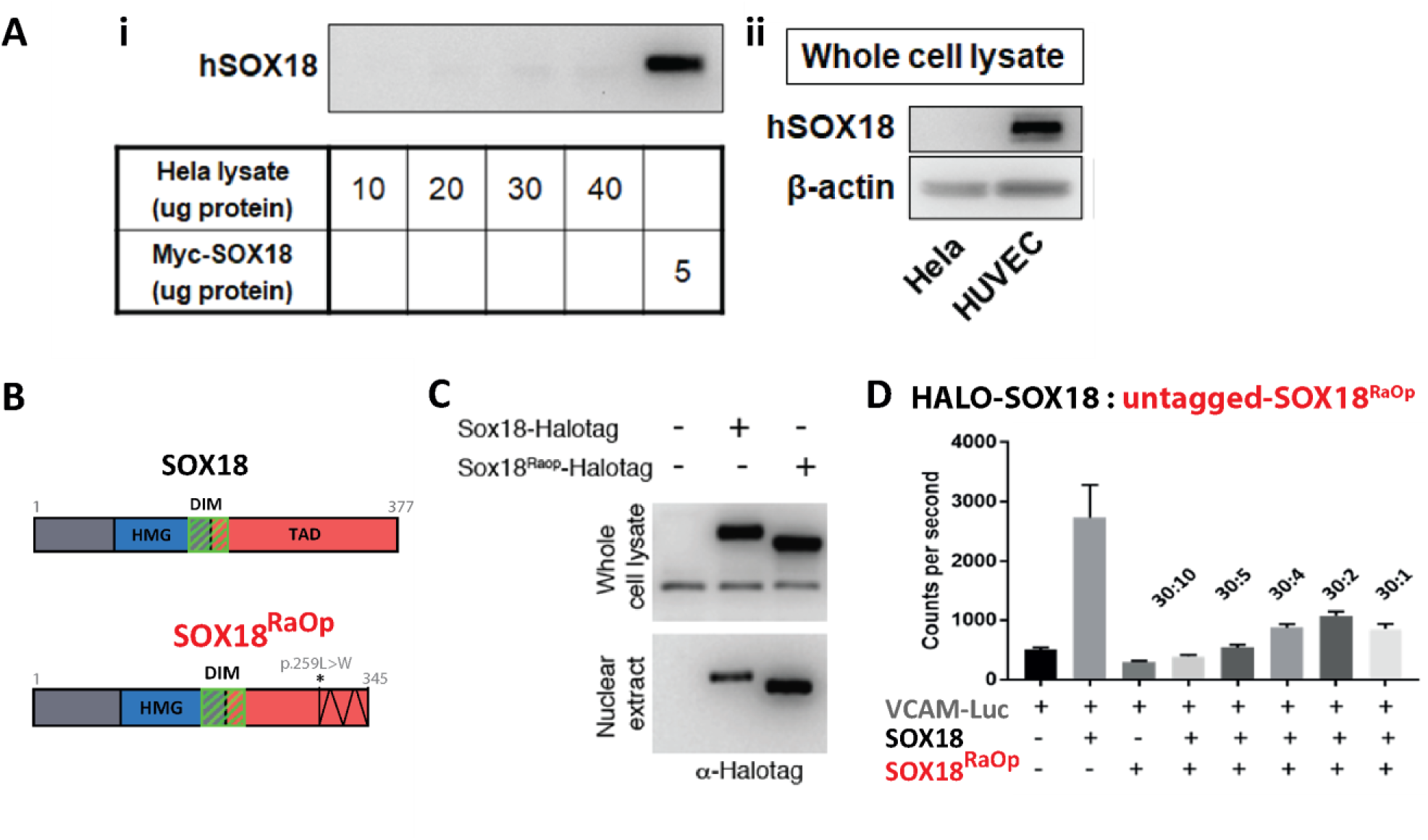
**HeLa cells show no detectable endogenous levels of SOX18 protein, the Ragged Opossum (SOX18^RaOp^) mutation does not interfere with SOX18 protein expression, and the HALO-tag does not interfere with the ability of SOX18 to activate gene transcription, or the ability of SOX18^RaOp^ to repress SOX18.** (A) Western blot analysis using an anti-human SOX18 antibody for human SOX18 (hSOX18) detection. (**i**) Lanes 1-4: 10, 20, 30 and 40 µg total protein from HeLa cell lysate. Lane 5: 5 µg of transfected human Myc-SOX18 protein. (**ii**) Western blot analysis on whole cell lysate from Hela cells (lane 1) and human umbilical venous endothelial cells (HUVECs; lane 2), with beta-actin used as a control. (**B**) Representation of SOX18 (top) and SOX18^RaOp^ (bottom) protein domains. SOX18 and SOX18^RaOp^ both share the same DNA-binding and bending HMG domain (blue) and homodimerization domain (DIM; green diagonal stripes), however SOX18^RaOp^ has a point mutation (c.775C>T; asterisk) in its C-terminal transactivation domain (red) which scrambles the rest of the domain (black zigzag) before resulting in a premature stop codon. (**C**) Western blot analysis on whole cell extracts from control conditions (empty vector) or transfected with HALO-SOX18 or HALO-SOX18^RaOp^ show a similar expression level of protein. Western-blot analysis from nuclear extracts (ctrl, HALO-SOX18 or HALO-SOX18^RaOp^) show an increase in the mutant protein. A halo-tag antibody was used for HALO-SOX18 or HALO-SOX18^RaOp^ detection. (**D**) Luciferase assay to measure VCAM1 promoter fragment transactivation (VCAM-Luc) in the presence of HALO-SOX18 and SOX18^RaOp^ protein. Validation of the dominant negative effect using different ratios of HALO-SOX18:SOX18^RaOp^ (30:10, 30:5, 30:4, 30:2 and 30:1). VCAM-Luc transactivation by SOX18 and SOX18^RaOp^ was performed in COS-7 cells, and is measured in counts per second, luciferase, arbitrary unit.

**Fig. S3 to Fig. 1.**
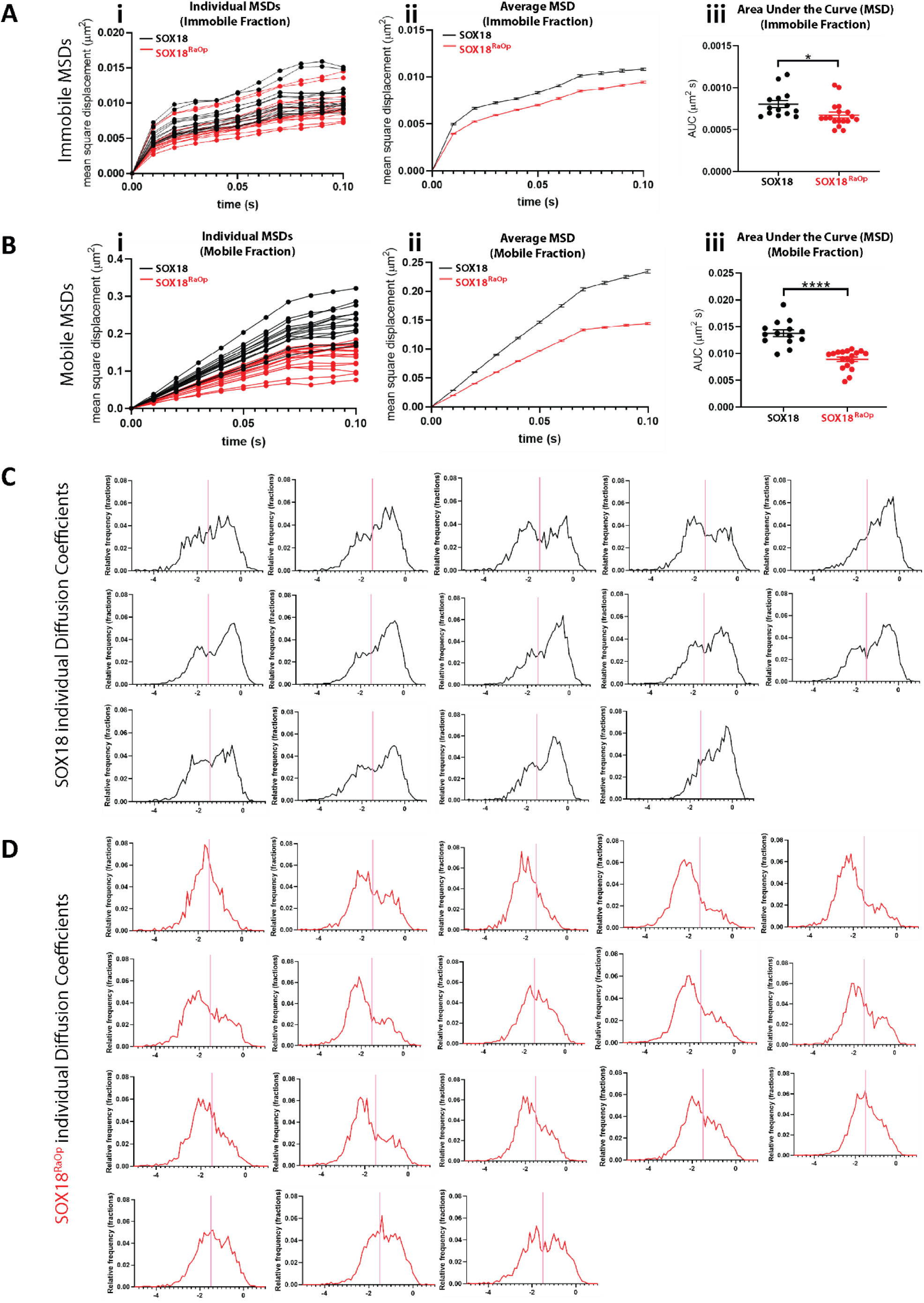
**Average MSDs with separated mobile and immobile fractions (top) and diffusion coefficient histograms (bottom) for each cell for HALO-SOX18 (black) and HALO-SOX18^RaOp^ (red).** (**A**) The average mean square displacement (MSD) for HALO-SOX18 (black) and HALO-SOX18^RaOp^ (red) separated into immobile fractions for each cell (**i**) and for each sample (**ii**). (**iii**) Quantification of the average MSD for immobile fractions for each cell is shown as area under the curve of the MSD. (**B**) The average mean square displacement (MSD) for HALO-SOX18 (black) and HALO-SOX18^RaOp^ (red) separated into mobile fractions for each cell **(i)** and for each sample (**ii**). (**iii**) Quantification of the average MSD for mobile fractions for each cell is shown as area under the curve of the MSD. (**C**) The average diffusion coefficient histogram for each cell for HALO-SOX18 (black), with the immobile to mobile threshold (Log_10_D = -1.5) shown by a pink line. (**D**) The average diffusion coefficient histogram for each cell for HALO-SOX18^RaOp^ (red), with the immobile to mobile threshold (Log_10_D = -1.5) shown by a pink line.

**Fig. S4 to Fig. 1.**
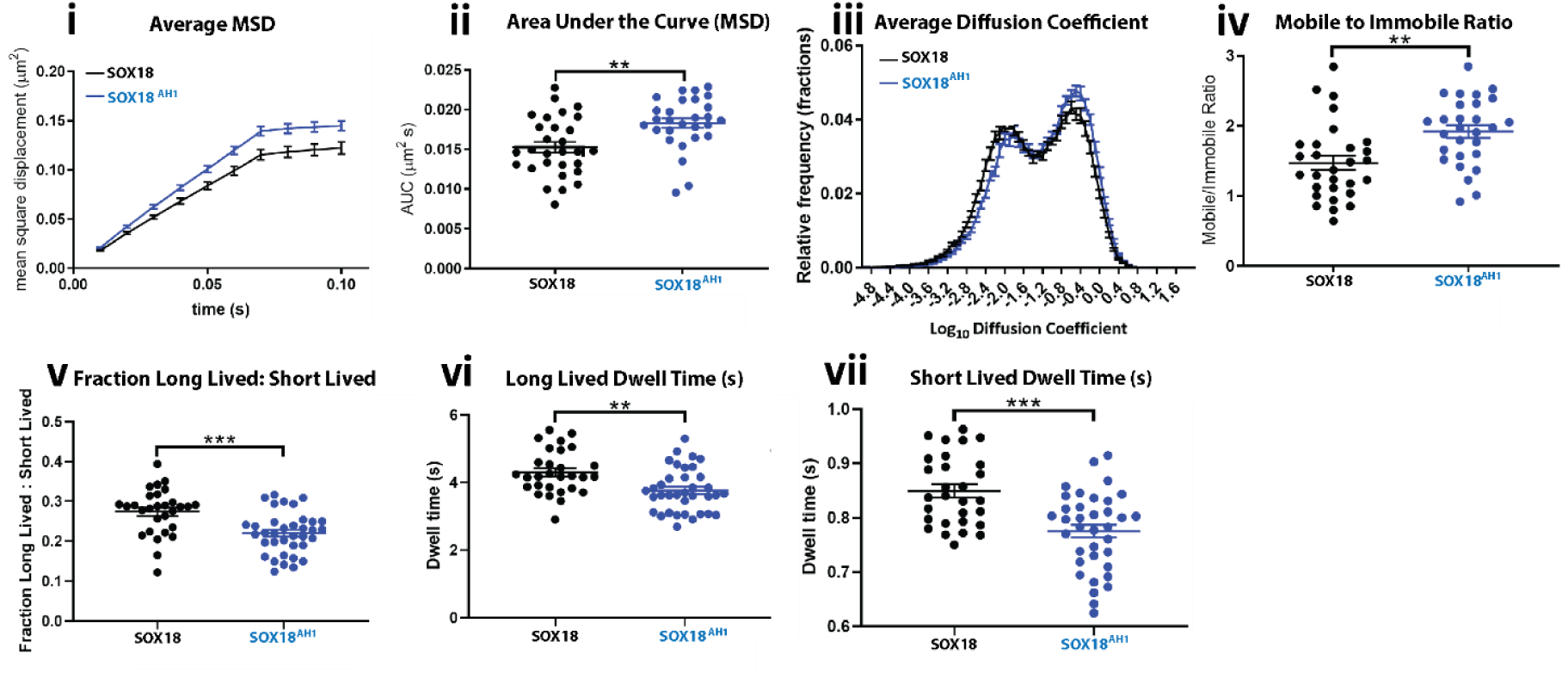
**A DNA-binding mutant (SOX18^AH1^) shows the opposite phenotype to SOX18^RaOp^.** Quantification of the dynamics of HALO-SOX18 (black) and HALO-SOX18^AH1^ (blue). **Top row:** quantification of fast tracking SMT data (20 ms acquisition for 6000 frames) represented by **(i)** the average mean square displacement (MSD; μm^2^s), **(ii)** the area under the curve of the average MSD for each cell (μm^2^s), **(iii)** the diffusion coefficient histogram for all cells (μm^2^s-1) and **(iv)** the mobile to immobile ratio for each cell. The threshold used to classify molecules as either mobile or immobile is Log_10_D = -1.5. n = 29 for HALO-SOX18 and n = 28 for HALO-SOX18^AH1^ (N = 3). t-test (two-tailed, unpaired). ns = non-significant. **Bottom row:** quantification of slow tracking SMT data (500 ms acquisition for 500 frames) showing **(v)** the fraction of long-lived to short-lived immobile events, and dwell times of **(vi)** long-lived and **(vii)** short-lived immobile events. Values for the mean ± s.e.m. are shown. Average number of trajectories obtained are 2713 for HALO-SOX18 and 1794 for HALO-SOX18^AH1^. n = 28 for HALO-SOX18 and n = 36 for HALO-SOX18^AH1^ (N = 3). Mann Whitney U-test for slow tracking (two-tailed, unpaired). ** P<0.01, **** P<0.0001.

**Fig. S1 to Fig. 3.**
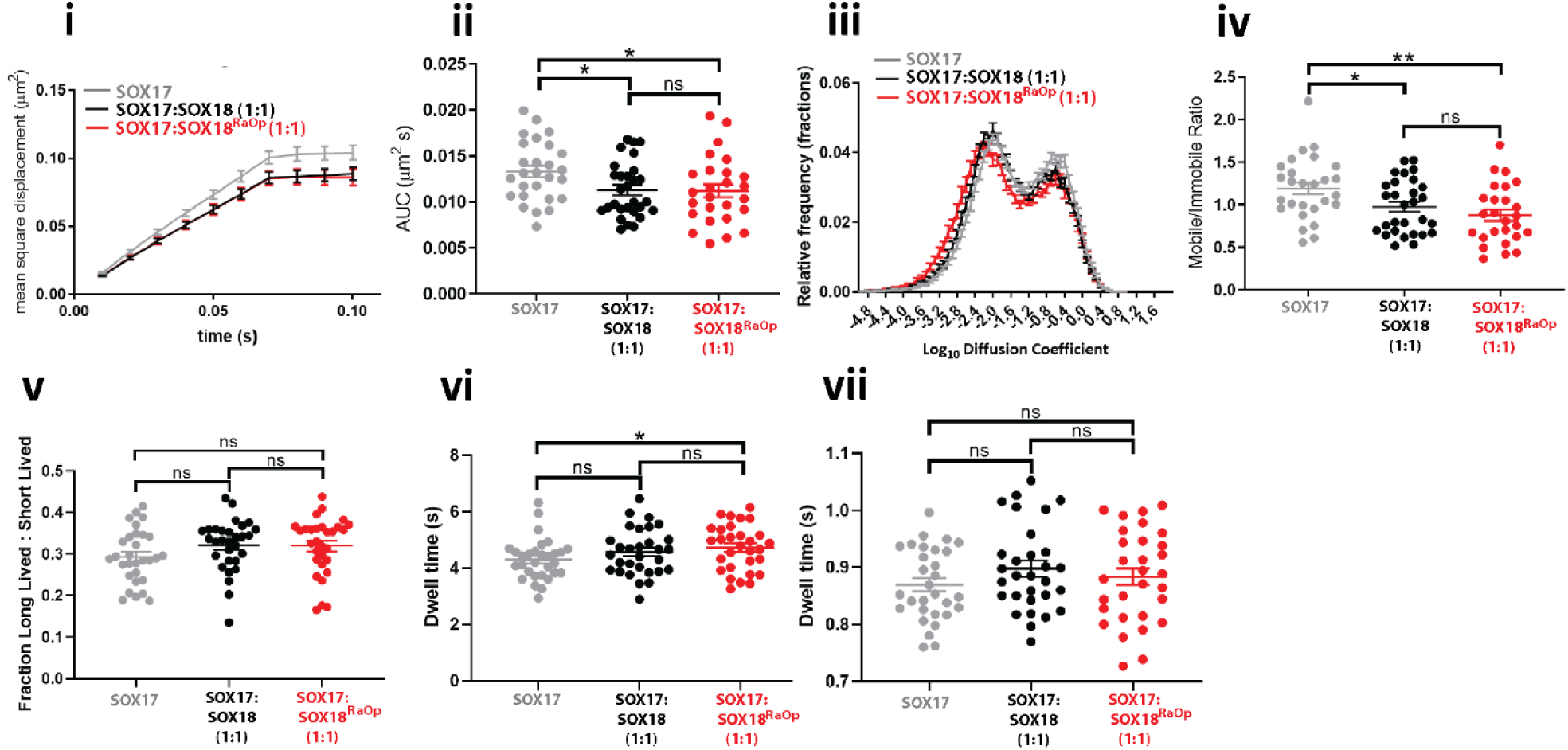
**SOX18^RaOp^ recruits SOXF member SOX17 in a similar fashion to wild-type SOX18.** Quantification of the dynamics of HALO-SOX17 (grey), HALO-SOX17 with untagged SOX18 in a 1:1 ratio (black) and HALO-SOX17 with untagged SOX18^RaOp^ in a 1:1 ratio (red). **Top row:** quantification of fast tracking SMT data (20 ms acquisition for 6000 frames) represented by (**i**) the average mean square displacement (MSD; μm^2^s), (**ii**) the area under the curve of the average MSD for each cell (μm^2^s), (**iii**) the diffusion coefficient histogram for all cells (μm^2^s-1) and (**iv**) the mobile to immobile ratio for each cell. The threshold used to classify molecules as either mobile or immobile is Log_10_D = -1.5. Values for the mean ± s.e.m. are shown. Average number of trajectories obtained are 1879 for HALO-SOX17, 1863 for HALO-SOX17:SOX18 (1:1) and 1676 for HALO-SOX17:SOX18^RaOp^ (1:1). n = 27 for HALO-SOX17, n = 29 for HALO-SOX17:SOX18 (1:1) and n = 26 for HALO-SOX17:SOX18^RaOp^ (1:1) (N = 3). t-test (two-tailed, unpaired). * P<0.05, ** P<0.01, ns = non-significant (P>0.05). **Bottom row:** quantification of slow tracking SMT data (500 ms acquisition for 500 frames) showing (**v**) the fraction of long-lived to short-lived immobile events and dwell times of (**vi**) long-lived and (**vii**) short-lived immobile events (s). Values for the mean ± s.e.m. are shown. n = 30 for HALO-SOX17, n = 30 for HALO-SOX17:SOX18 (1:1) and n = 30 for HALO-SOX17:SOX18^RaOp^ (1:1) (N = 3). Mann Whitney U-test for slow tracking (two-tailed, unpaired). * P<0.05, ** P<0.01, ns = non-significant (P>0.05).

**Fig. S1 to Fig. 4.**
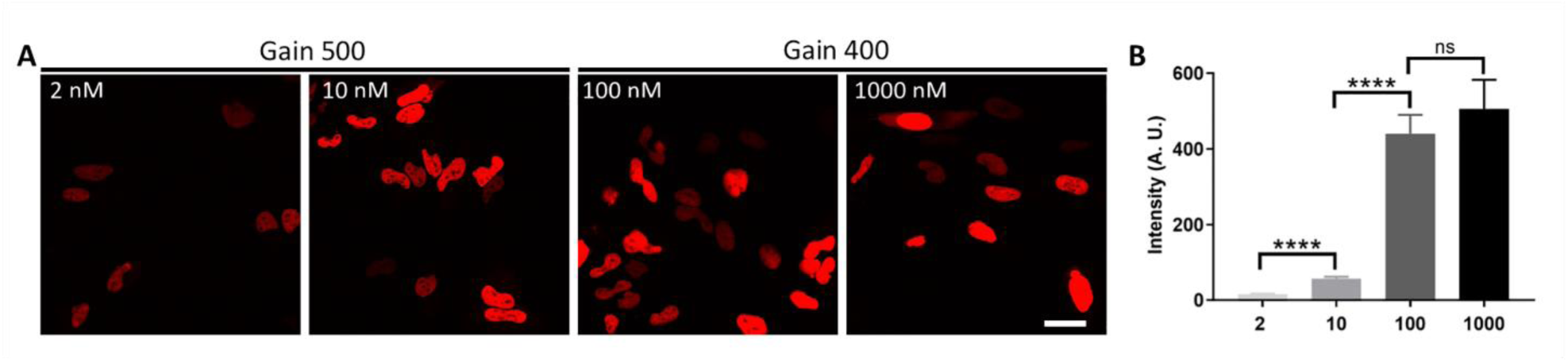
**JF549 dye titration to identify an optimum concentration for saturating HALO-SOX18 molecules for use in N&B and cRICS experiments.** (**A**) Confocal images showing HeLa cells transfected with HALO-SOX18 and exposed to a titration of JF549 dye. (**B**) Titration of JF549 dye shows that HALO-SOX18 becomes saturated with dye at approximately 1000 nM, with no significant change in intensity observed after 100 nM.

**Fig. S2 to Fig. 4.**
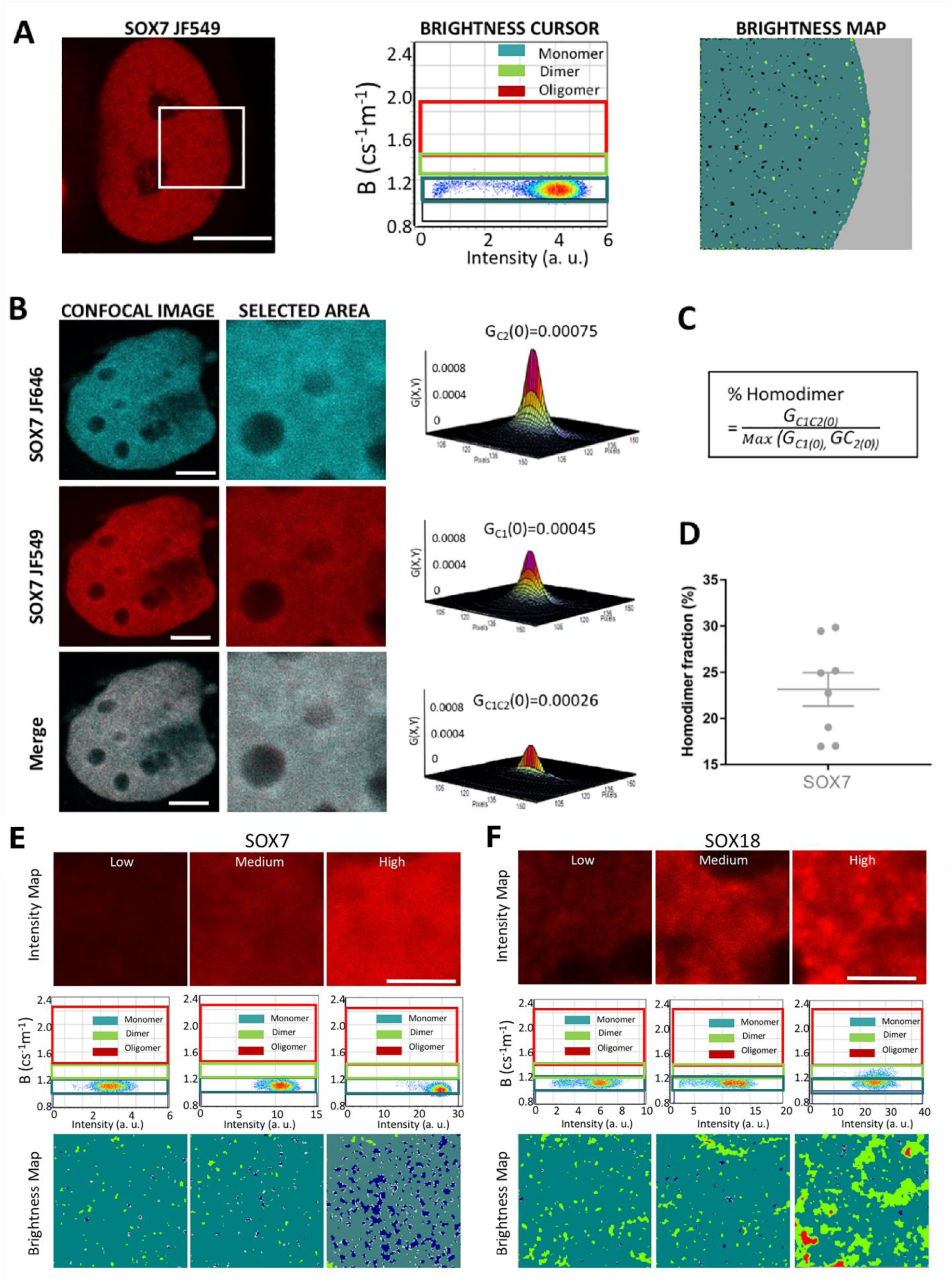
**SOX7 monomer brightness is used to calibrate number and brightness (N&B) and cross-raster image correlation spectroscopy (cRICS) analyses.** (A) SOX7 does not form homodimers and therefore has been used as a control to calibrate the brightness for single HALO-JF549 molecules in number and brightness (N&B) analyses. **(Left)** HeLa cells are transfected with HALO-SOX7 and exposed to 1 µM of JF549 dye. The selection area on which N&B is performed is outlined by a white square. Scale bar = 10 µM. **(Middle)** The brightness cursor showing the separation of SOX7 into different oligomeric states based on brightness. As SOX7 is monomeric, the monomer cursor (dark green) for all experiments is set based on SOX7 brightness. The homodimer cursor (light green) is of the same width and placed directly above the monomer cursor, and the higher order oligomer (three or more molecules in a complex) is placed so that it contains all brightness values above the homodimer cursor. An absence of molecules is shown by a dark blue cursor directly below the monomer cursor. **(Right)** Brightness maps indicate the oligomeric distribution of HALO-SOX7, with monomers/heterodimers being represented as dark green pixels, “homodimers” as light green pixels, and an absence of molecules as dark blue pixels. (**B**) **(Left column):** HeLa cells were transfected with HALO-SOX7 and exposed to 500 µM of JF549 and JF646 HaloTag dye. (Scale bar = 5µM). The JF646 excitation channel is shown on the top row, the JF549 excitation channel is shown in the middle row, and the merging of these two channels is shown on the bottom row. The section area of the nucleus on which cRICS analysis was performed is outlined by a white box. **(Middle column):** The selection area on which N&B analysis was performed. **(Right column):** The 3-dimentional cRICS functions for the individual channels are shown, as well as the cross-correlation between these channels. (**C**) The equation used to derive the percentage of homodimers using cRICS. (**D**) Quantification of the percentage of SOX7 molecules in juxtaposition in **C,** using the equation in **D,** as a control to calibrate the false positive cross-correlation due to the spectrum bleed through. (HALO-SOX7 n = 8). Values for the mean ± s.e.m. are shown. (**E**) N&B analysis performed on HALO-SOX7 using selected HeLa cells with different HALO-SOX7 concentrations. **(Top row)** An example nucleus of a cell with low HALO-SOX7 expression is shown on the left, medium expression in the middle, and high expression on the right. **(Middle row)** The brightness cursor showing the separation of HALO-SOX7 into different oligomeric states based on brightness. As SOX7 is monomeric, the monomer cursor (dark green) is set based on HALO-SOX7 brightness. The homodimer cursor (light green) is of the same width and placed directly above the monomer cursor, and the higher order oligomer (three or more molecules in a complex) is placed so that it contains all brightness values above the homodimer cursor. An absence of molecules is shown by a dark blue cursor directly below the monomer cursor. **(Bottom row)** Brightness maps indicate the oligomeric distribution of HALO-SOX7, with monomers/heterodimers being represented as dark green pixels, homodimers as light green pixels and an absence of molecules as dark blue pixels. (**F**) N&B analysis performed on HALO-SOX18 using selected HeLa cells with different HALO-SOX18 concentrations. **(Top row)** An example nucleus of a cell with low HALO-SOX7 expression is shown on the left, medium expression in the middle, and high expression on the right. **(Middle row)** The brightness cursor showing the separation of SOX18 into different oligomeric states based on brightness. The cursors are set based on the brightness values obtained for the HALO-SOX7 monomeric control. The monomer cursors are shown in dark green, the homodimer cursor in light green, the higher order oligomer cursor in red, and an absence of molecules in dark blue. **(Bottom row)** Brightness maps indicate the oligomeric distribution of HALO-SOX18, with monomers/heterodimers being represented as dark green pixels, homodimers as light green pixels, higher-order oligomers as red pixels and an absence of molecules as dark blue pixels. (Scale bar = 5 µm).

**Fig. S3 to Fig. 4.**
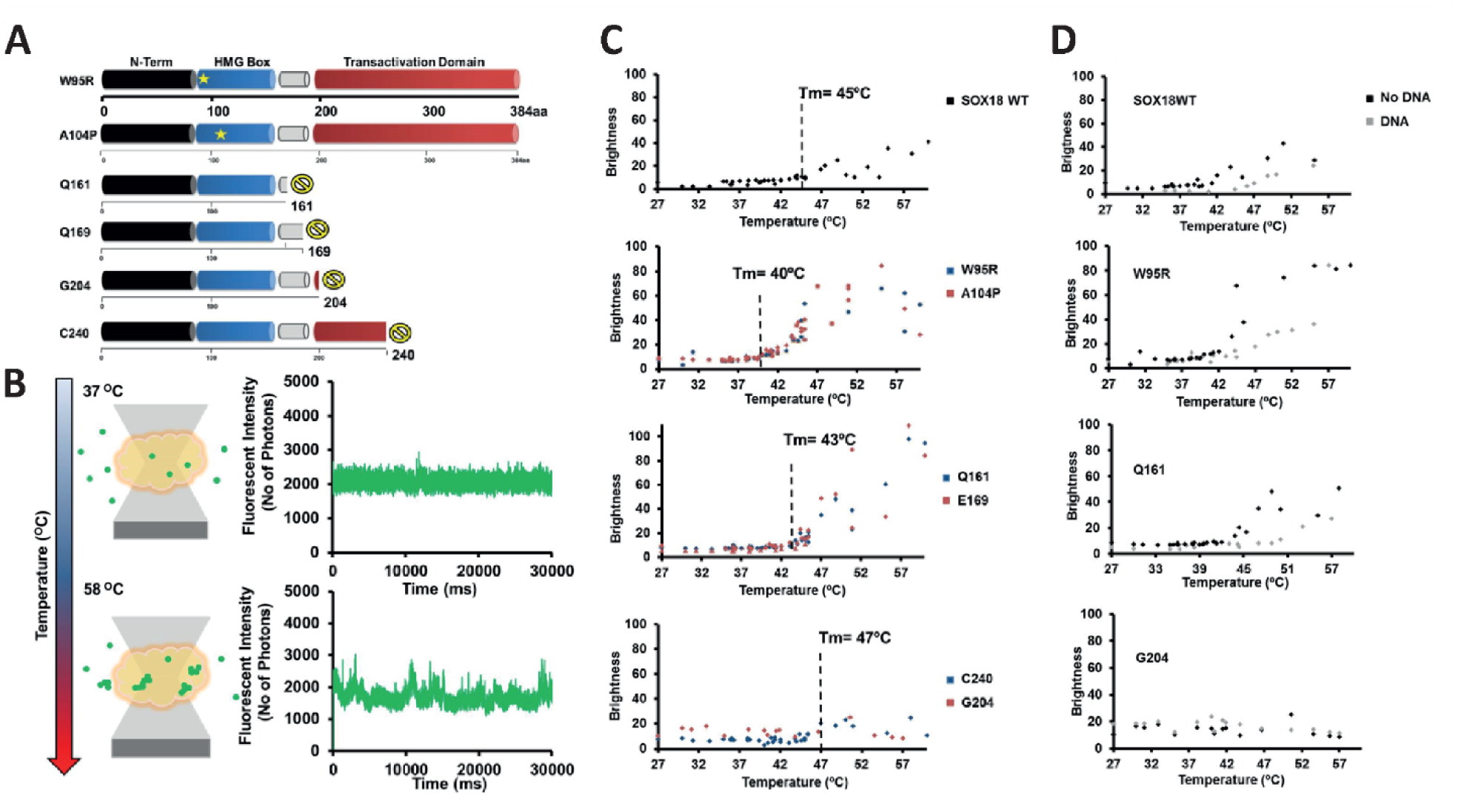
**Different mutations within SOX18 alters its likelihood to form homodimers and its chromatin-binding stability.** (A) Schematic showing the protein domains of SOX18 and various recessive (W95R and A104P) and dominant-negative (Q161*, Q169*, G204* and C240*) mutations reported to cause Hypotrichosis-lymphedema-telangiectasia-renal Syndrome (HLTRS) in humans. A star indicates a point mutation, and a yellow ‘no’ symbol denotes premature truncations. SOX18 contains an N-terminal domain (black), followed by a DNA-binding and bending HMG domain (blue), a homodimerisation domain (white) and a C-terminal transactivation domain (red). (**B**) Schematic showing the fluorescence fluctuation single molecule assay performed to identify the degree of SOX18 aggregation at different temperatures (37 °C and 58 °C) which assesses the level of protein stability. Aggregation is shown by an increase in the fluctuation of fluorescence intensity indicating that multiple proteins are passing through together (bottom). Proteins with higher stabilities will require higher temperatures before they become aggregated. (**C**) Brightness assay assessing protein stability of SOX18 and various SOX18 mutants determined by the temperature at which they become aggregated (dotted line). (**D**) Brightness assay assessing protein stability of SOX18 and various SOX18 mutants in the presence (grey) and absence of DNA (black).

**Fig. S4 to Fig. 4.**
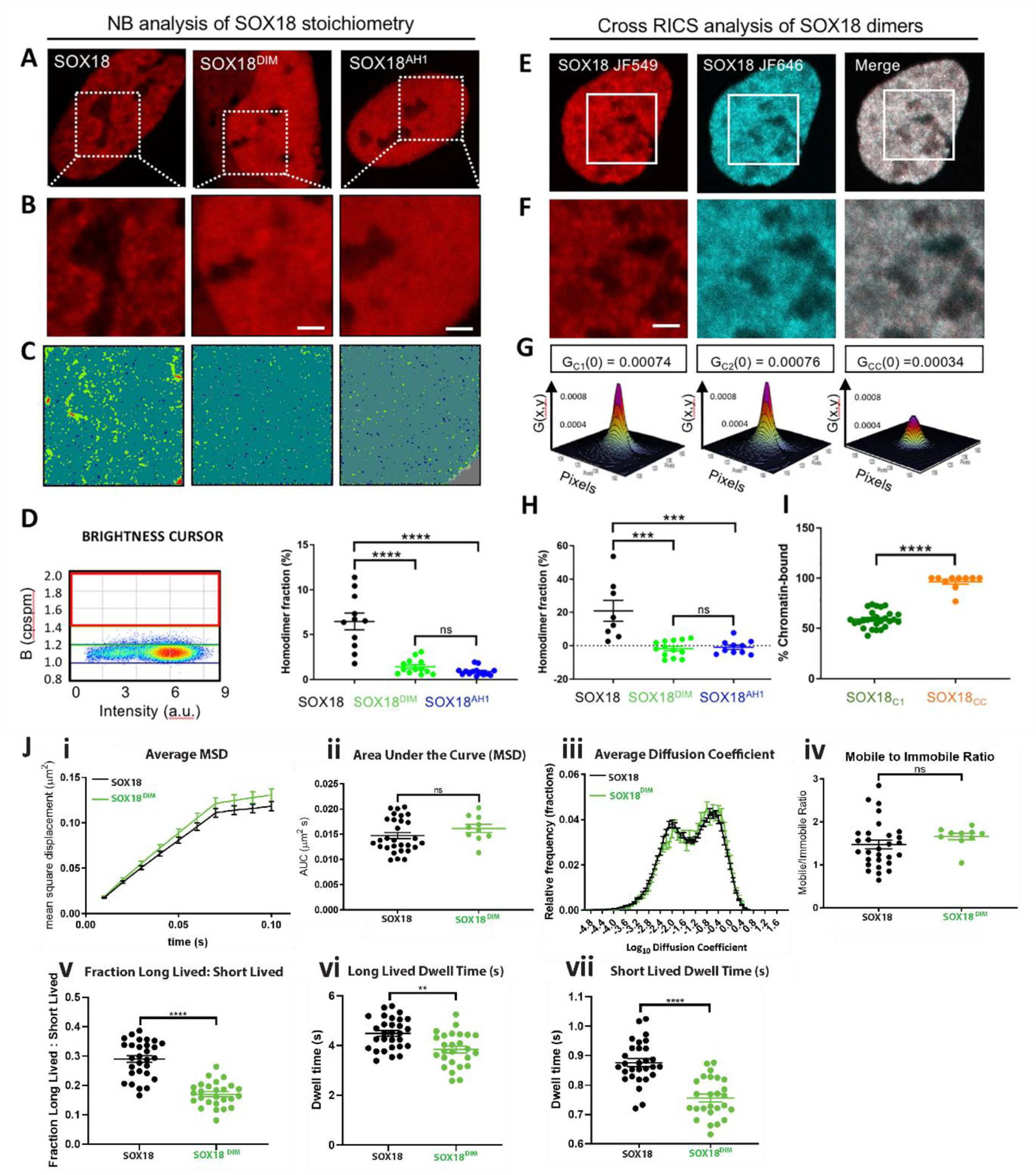
**SOX18 forms homodimers on the chromatin via a DNA-dependent cooperative mechanism.** (**A-D**) Number and brightness (N&B) analysis of the oligomeric distribution of HALO-SOX18, HALO-SOX18^AH1^ and HALO-SOX18^DIM^ in the nucleus of HeLa cells. (**A**) HeLa cells are transfected with HALO-SOX18 (left), HALO-SOX18^DIM^ (middle) or HALO-SOX18^AH1^ (right) and exposed to 2 nM of JF549 HaloTag dye. The section area of the nucleus on which N&B analysis was performed is outlined by a white box. (**B**) The selection area on which N&B analysis was performed. Scale bar = 2 µM. (**C**) The brightness map showing the distribution of SOX18 monomers (dark green), homodimers (light green) and higher order oligomers (red). An absence of tagged molecules is shown in dark blue. (**D**) (**Left**) Intensity versus brightness scatterplot of the N&B data acquisition presented for HALO-SOX18. SOX18 monomers are outlined by a dark green selection box, SOX18 homodimers by a light green selection box and SOX18 higher order oligomers (3 or more SOX18 molecules in a complex) by a red selection box. The position of the selection boxes was determined based on a HALO-SOX7 monomeric control (shown in Fig. S2A-E to Fig. 4). (Right) The percentage of homodimers for HALO-SOX18 (black), HALO-SOX18DIM (green) and HALO-SOX18AH1 (blue). Values for the mean ± s.e.m. are shown. HALO-SOX18 n = 11, HALO-SOX18DIM n = 13 and HALO-SOX18AH1 n = 14. Mann Whitney U-test for slow tracking (two-tailed, unpaired). **** P<0.0001, ns = non-significant. (**E-H**) Cross-raster image correlation spectroscopy (cRICS) analysis to validate the formation of HALO-SOX18, HALO-SOX18DIM and HALO-SOX18AH1 homodimers. (**E**) HeLa cells were transfected with HALO-SOX18 and exposed to 500 nM of JF549 and JF646 HaloTag dye. The JF549 excitation channel is shown on the left, the JF646 excitation channel is shown in the middle, and the merging of these two channels is shown on the right. The section area of the nucleus on which cRICS analysis was performed is outlined by a white box. (**F**) The selection area on which cRICS analysis was performed. Scale bar = 2 µM. (**G**) The 3D cRICS functions obtained for HALO-SOX18 in the JF549 channel (left), the JF646 channel (middle) and the cross-correlation of these two channels (right). (**H**) The fraction of homodimers/oligomers for the HALO-SOX18, HALO-SOX18^DIM^ and HALO-SOX18^AH1^. Values for the mean ± s.e.m. are shown. HALO-SOX18 n = 8, HALO-SOX18^DIM^ n = 13 and HALO-SOX18^AH1^ n = 10. Statistical significance was assessed using Mann Whitney U-test for slow tracking (two-tailed, unpaired). *** P<0.001, ns = non-significant. (**I**) The fraction of chromatin-bound HALO-SOX18 homodimers/oligomers. The percentage of homodimers identified in a single channel (homodimers with the same JF549 dye) is shown in green, and cross-correlation of both channels to identify the percentage of homodimers with two dyes (homodimers with 1 JF549 dye and 1 JF646 dye) is shown in orange. Values for the mean ± s.e.m. are shown. HALO-SOX18 homodimers in a single channel (orange) had a sample size of n = 27 and HALO-SOX18 correlated across both channels (green) had a sample size of n = 10. Mann Whitney U-test for slow tracking (two-tailed, unpaired). **** P<0.0001. (**J**) Quantification of the dynamics of HALO-SOX18 (black) and HALO-SOX18^DIM^ (green). **Top row:** quantification of fast tracking SMT data (20 ms acquisition for 6000 frames) represented by **(i)** the average mean square displacement (MSD; μm^2^s), **(ii)** the area under the curve of the average MSD for each cell (μm^2^s), **(iii)** the diffusion coefficient histogram for all cells (μm^2^s-1) and **(iv)** the mobile to immobile ratio for each cell. The threshold used to classify molecules as either mobile or immobile is Log_10_D = -1.5. n = 28 for HALO-SOX18 and n = 10 for HALO-SOX18^DIM^ (N = 3). t-test (two-tailed, unpaired). ns = non-significant. **Bottom row:** quantification of slow tracking SMT data (500 ms acquisition for 500 frames) showing **(v)** the fraction of long-lived to short-lived immobile events, and dwell times of **(vi)** long-lived and **(vii)** short-lived immobile events. Values for the mean ± s.e.m. are shown. Average number of trajectories obtained are 2379 for HALO-SOX18 and 1356 for HALO-SOX18^DIM^. n = 29 for HALO-SOX18 and n = 26 for HALO-SOX18^DIM^ (N = 3). Mann Whitney U-test for slow tracking (two-tailed, unpaired). ** P<0.01, **** P<0.0001.

**Fig. S5 to Fig. 4.**
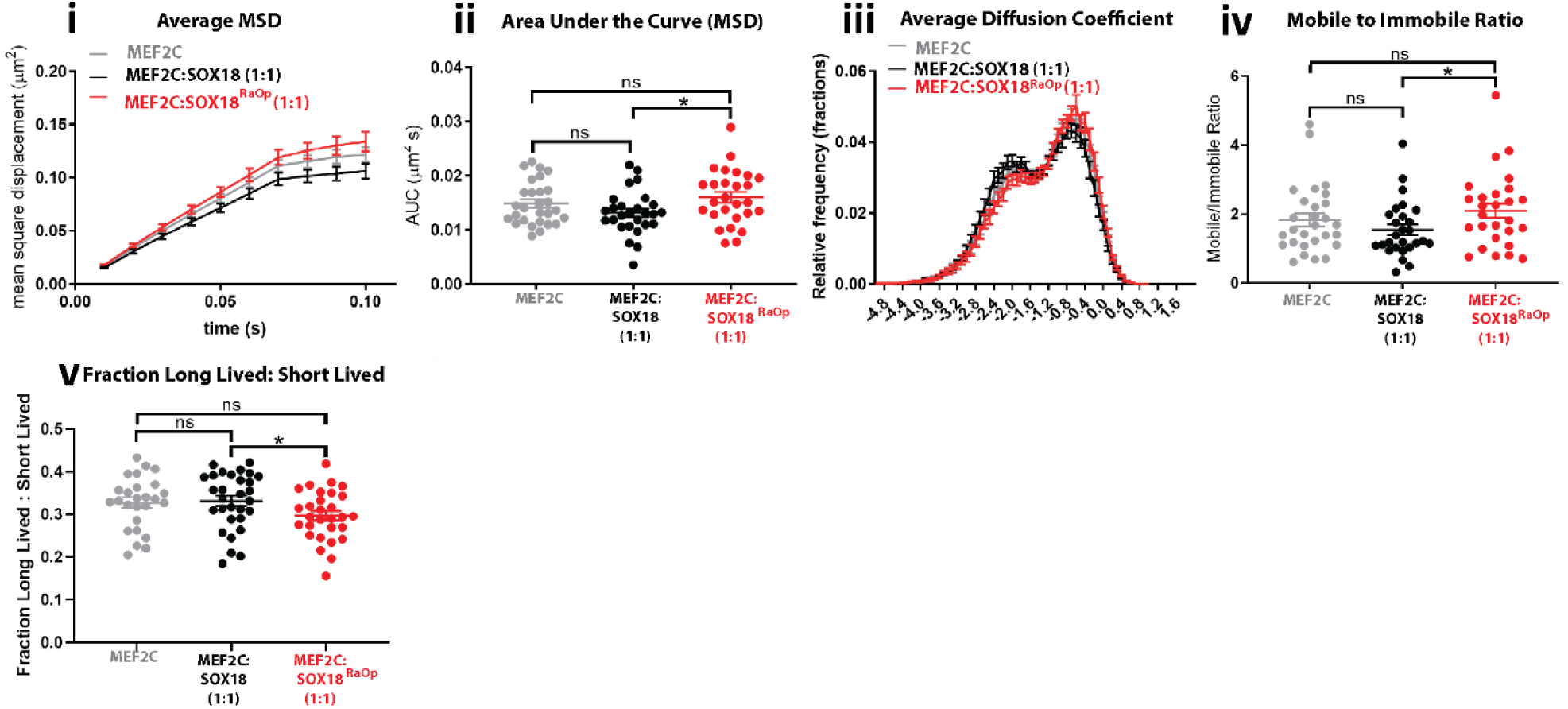
**SOX18^RaOp^ causes an increase in MEF2C diffusion and a decrease in MEF2C chromatin-bound and long-lived binding fractions compared to SOX18.** Quantification of the dynamics of HALO-MEF2C (grey), HALO-MEF2C with untagged SOX18 in a 1:1 ratio (black) and HALO-MEF2C with untagged SOX18^RaOp^ in a 1:1 ratio (red). **(Top row):** quantification of fast tracking SMT data (20 ms acquisition for 6000 frames) represented by (**i**) the average mean square displacement (MSD; μm^2^s), (**ii**) the area under the curve of the average MSD for each cell (μm^2^s), (**iii**) the diffusion coefficient histogram for all cells (μm^2^s-1) and (**iv**) the mobile to immobile ratio for each cell. The threshold used to classify molecules as either mobile or immobile is Log_10_D = -1.5. Values for the mean ± s.e.m. are shown. The average number of trajectories obtained are 2275 for HALO-MEF2C, 2408 for HALO-MEF2C:SOX18 (1:1) and 1737 for HALO-MEF2C:SOX18^RaOp^ (1:1). n = 28 for HALO-MEF2C, n = 27 for HALO-MEF2C:SOX18 (1:1) and n = 27 for HALO-MEF2C:SOX18^RaOp^ (1:1) (N = 3). t-test (two-tailed, unpaired). * P<0.05, ns = non-significant (P>0.05). **(Bottom row):** quantification of slow tracking SMT data (500 ms acquisition for 500 frames) showing **(v)** the fraction of long-lived to short-lived immobile events. Values for the mean ± s.e.m. are shown. n = 26 for HALO-MEF2C, n = 30 for HALO-MEF2C:SOX18 (1:1) and n = 29 for HALO-MEF2C:SOX18^RaOp^ (1:1) (N = 3). Whitney U-test for slow tracking (two-tailed, unpaired). * P<0.05, ns = non-significant (P>0.05).

### Supplemental Tables

**Table S1:**
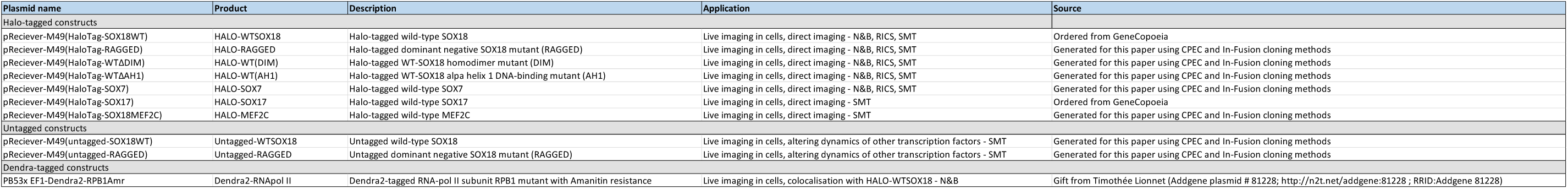
**Plasmid information.**

## Supplemental Videos

Examples of single molecule tracking movies acquired for each condition using fast and slow tracking methods can be accessed using the links below:

Video S1 – SOX18 VS SOX18RaOp (FAST TRACKING) Related to Fig. 1. https://cloudstor.aarnet.edu.au/plus/s/KtM31tcucdqxdCq

Video S2 – SOX18 VS SOX18RaOp (SLOW TRACKING). Related to Fig. 1. https://cloudstor.aarnet.edu.au/plus/s/r4j0sBeEYCjOnXF

Video S3 – SOX18 WITH SOX18RaOp (FAST TRACKING). Related to Fig. 2. https://cloudstor.aarnet.edu.au/plus/s/mpc3G9md7Rpq5hV

Video S4 – SOX18 WITH SOX18RaOp (SLOW TRACKING). Related to Fig. 2. https://cloudstor.aarnet.edu.au/plus/s/BCKpYDzkIXN6Ywd

Video S5 – SOX7 (FAST TRACKING). Related to Fig. 3. https://cloudstor.aarnet.edu.au/plus/s/nvyLNL8BlNLb6SX

Video S6 – SOX7 (SLOW TRACKING). Related to Fig. 3. https://cloudstor.aarnet.edu.au/plus/s/zrp7SfWzTMQ0mxk

Video S7 – SOX17 (FAST TRACKING). Related to Fig. S1 to Fig. 3. https://cloudstor.aarnet.edu.au/plus/s/8t1vFrdL8GJQzaJ

Video S8 – SOX17 (SLOW TRACKING). Related to Fig. S1 to Fig. 3. https://cloudstor.aarnet.edu.au/plus/s/Cks0ZxLCGbohiWA

Video S9 – SOX18DIM (FAST TRACKING). Related to Fig. S3 to Fig. 4. https://cloudstor.aarnet.edu.au/plus/s/hx5Kzl86VjAiyvj

Video S10 – SOX18DIM (SLOW TRACKING). Related to Fig. S3 to Fig. 4. https://cloudstor.aarnet.edu.au/plus/s/Wx5kyt0f9YTTZgm

Video S11 – SOX18AH1 (FAST TRACKING). Related to Fig. S3 to Fig. 4. https://cloudstor.aarnet.edu.au/plus/s/MMtMkTgjSucUlJr

Video S12 – SOX18AH1 (SLOW TRACKING). Related to Fig. S3 to Fig. 4. https://cloudstor.aarnet.edu.au/plus/s/G8xjmozoWOGkLry

Video S13 – MEF2C (SLOW TRACKING) Related to Fig. 4 and Fig. S5 to Fig. 4. https://cloudstor.aarnet.edu.au/plus/s/SbAJpLL0cOrZyJX

Video S14 – MEF2C (FAST TRACKING). Related to Fig. S5 to Fig. 4. https://cloudstor.aarnet.edu.au/plus/s/zER5kfUMv9A9WWV

